# Unveiling Novel Hyperglycosylated Analog of Human Erythropoietin: A Comprehensive Computational Exploration

**DOI:** 10.1101/2024.10.07.617028

**Authors:** Naima Thahsin, Khairul Islam Khan, Mohammad Nazmus Sakib, Mohammad Shahedur Rahman

## Abstract

**Introduction:** The capability of human erythropoietin (hEPO) to treat anemia by stimulating erythropoiesis, has made it become one of the world’s leading biopharmaceuticals. An effective way to improve the efficiency of recombinant hEPO drugs is increasing glycosylation which can increase its serum half-life and thereby it’s *in vivo* biological activity. In this context, the objective of the present study was to explore potential glycosylation sites in hEPO to develop new hyperglycosylated analogous drug.

**Materials and Methods:** A rational computational strategy was executed to select compatible region of hEPO for inserting N-linked glycosylation consensus motif (N-X-S/T), thus designing a number of analogs. The 3D models of the analogs were constructed by homology modeling and validated by various *in silico* tools. The probability of glycosylation was checked and hyperglycosylated models of the selected analogs were developed. Molecular docking and molecular dynamics simulation were performed to select the best analog.

**Results:** Among 40 analogs designed in this study, Analog-71.1 showed most promising results. Four possible sites of this analog showed good probability for glycosylation. According to the molecular docking study, the analog obtained high docking score (2313.711) and formed highest number of (25) hydrogen bonds with erythropoietin receptors, which indicated strong and stable interaction with the receptors. Moreover, the steady trajectory of RMSD, R_gyr_, RMSF and other findings obtained from MD simulation confirmed the structural stability of the analog.

**Conclusions:** Our overall study has selected Analog-71.1 as a potential candidate to develop new analogous drug with enhanced efficiency, which needs further experimental analyses.

## Introduction

Human erythropoietin (hEPO) is an essential hematopoietic growth factor which regulates erythropoiesis by stimulating the proliferation and differentiation of erythroid progenitor cells in the bone marrow and protecting them from cellular apoptosis (Thomson and Lotze, 2003). Erythropoietin is synthesized as a 193 amino acid long prohormone which becomes 165 amino acid long mature hormone after the cleavage of the leader sequence (27 amino acids) and the arginine residue from the N and C terminals, respectively (Lai et al., 1986). It is a heavily glycosylated protein which has three N-linked sugars attached to the amine groups of asparagines at positions 24, 38 and 83 and one O-linked sugar attached to the hydroxyl group of serine at position 126 (Browne et al., 1986; Egrie et al., 1986; Sasaki et al., 1987; Tsuda et al., 1988). The carbohydrate structures of hEPO exhibit the feature of microheterogeneity which articulates the fact that, although the amino acid sequence of protein portion of the hormone is invariant, the oligosaccharides attached to it are highly variable in structure. For example, the N-linked saccharides of hEPO can be biantennary, triantennary, triantennary with one N-acetyllactosaminyl repeat, tetraantennary and tetraantennary with one, two, or three N-acetyllactosaminyl repeats (Sasaki et al., 1987). The variability in hEPO carbohydrates is further detected in the number of sialic acids which are the typical terminal molecules of these carbohydrate chains. The total number of sialic acid residues on one hEPO molecule can be as low as 0 to as high as 14 (Egrie & Browne, 2001).

In adult human body the journey of hEPO starts predominantly from kidney in response to tissue oxygenation level (Jacobson et al., 1957), which then travels through the circulatory system to the bone marrow and binds to the erythropoietin receptors (EPORs) expressed on the cell surface of red blood cell precursors (Sawada et al., 1990). The interaction between hEPO and EPORs leads to receptor dimerization (Philo et al., 1996), along with some conformational changes in the receptors that activates the cytoplasmic tyrosine kinase, JAK2, which remains pre-bound to the cytoplasmic tail of the receptors (Witthuhn et al., 1993). The JAK2, in turn, phosphorylates several cytoplasmic tyrosine residues of the EPORs that act as the docking sites for a number of signaling molecules (Richmond et al., 2005; Thomson & Lotze, 2003). The activation of all these signaling molecules transmits necessary signals for the expression of genes and synthesis of biomolecules essential for the survival, multiplication and maturation of the erythroid progenitor cells, consequently increasing the red blood cell mass. The role of erythropoietin in increasing the hematocrit has made it become one of the world’s leading biopharmaceuticals. Since the end of 1980s, recombinant human erythropoietin (rhEPO) has been granted as a therapeutic agent for the treatment of anemia and produced commercially (Eschbach et al., 1987). The rhEPO drug has been extremely helpful in various clinical conditions associated with anemia like chronic renal failure, HIV infection, cancer and even chemotherapy (Elliott et al., 2003). Due to the rapid clearance from the serum, the rhEPO has to be administered frequently to the anemic patients which is two to three times per week (Elliott et al., 2003). Any modification in the bioactivity of rhEPO by increasing its serum half-life would require less frequent administration and thus be advantageous for patients. Several attempts have been made to boost up the bioactivity of the recombinant hEPO drug, one of which was to manipulate its post-translational modification, particularly glycosylation.

The glycosylation has very vital impact upon the biological function of erythropoietin (Higuchi et al., 1992). It has been found that increasing the quantity of sialic acid containing glycan moieties of the hormone, increases its serum half-life, thereby prolonging its duration of action in regulating erythropoiesis (Egrie & Browne, 2002). The glycoengineered analog of rhEPO (Darbepoetin alpha) containing two additional N-linked carbohydrate chains at positions 30 and 88, has shown enhanced bioactivity (Egrie & Browne, 2001, 2002). Other potential positions of the hormone for glycosylation should be evaluated to produce biosimilar drugs. However, constructing multiple analogs of a protein in laboratory is usually time consuming and expensive. Using computational methods to design and model analogs and predict their efficiency prior to laboratory work would be helpful. In this context, the main objective of the current study was to design and model hyperglycosylated hEPO analogs with additional glycosylation through *in silico* tools and select the best analog on the basis of its stable structural conformation and better interaction with the receptors.

## Materials and Methods

The workflow used in this study is shown step by step in Figure 1.

**Figure 1:**
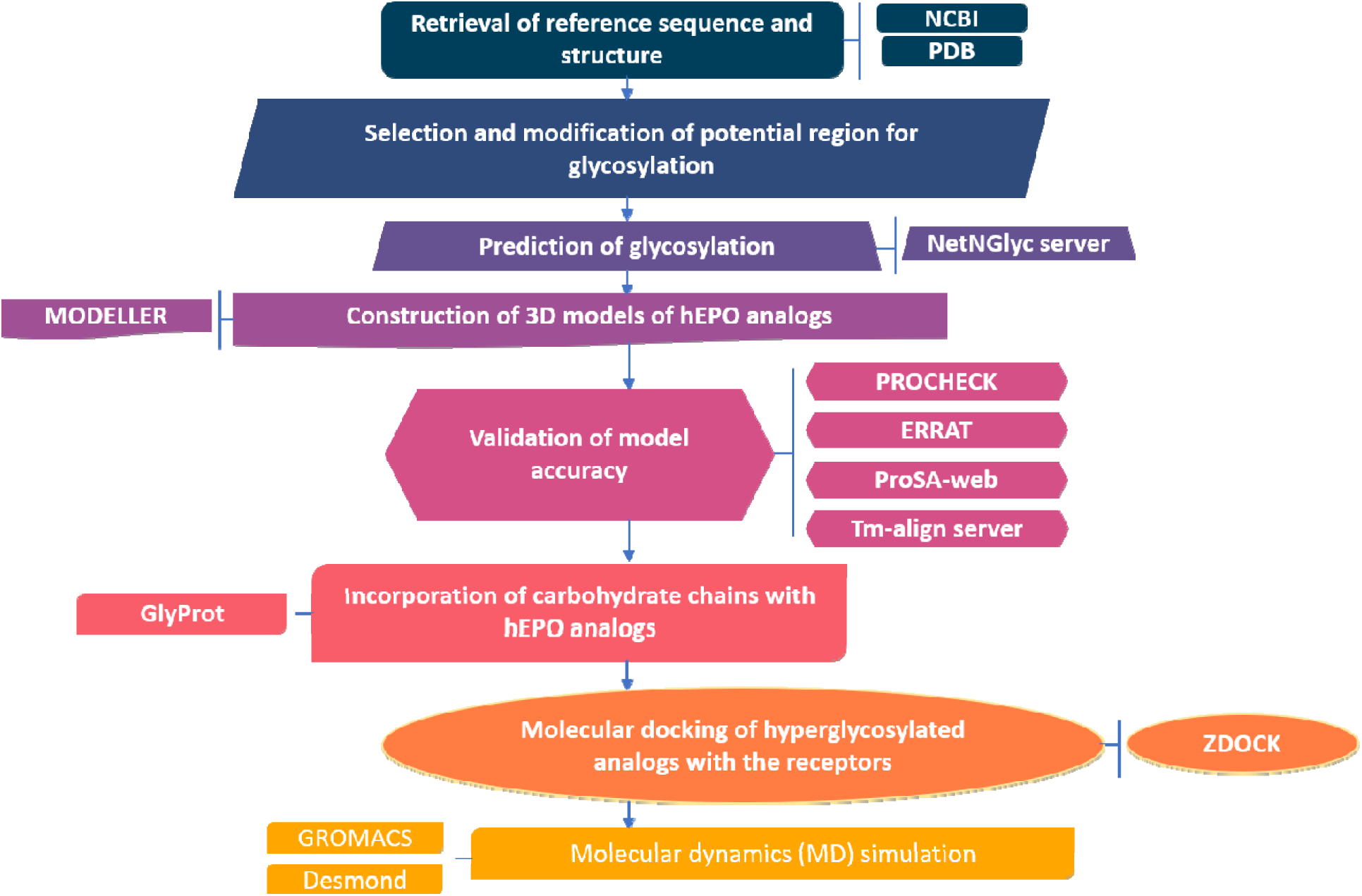
Schematic illustration of the study’s workflow. The procedure involves exploration of potential glycosylation sites in hEPO to develop new hyperglycosylated analogous drug.

### Retrieval of reference sequence and structure

The amino acid sequence of human erythropoietin (hEPO) was retrieved in FASTA format from the NCBI database (accession ID: AAA52400) (Lin et al., 1985) (https://www.ncbi.nlm.nih.gov/) to be used as the reference sequence in this study. The retrieved sequence was a prepeptide containing 193 amino acids among which 1-27 residues correspond to a signal peptide which was removed before modification of the sequence and modeling of analogs. To use as the reference structure, the three-dimensional (3D) structure of the hEPO molecule was extracted from the RCSB Protein Data Bank (PDB ID: 1EER) (Berman et al., 2000) (http://www.rcsb.org/pdb/). It is a complex crystal structure (resolution 1.9 Å) of three molecules containing one erythropoietin molecule and two erythropoietin receptors.

### Selection and modification of potential region for glycosylation

To increase the carbohydrate content of human erythropoietin, additional consensus motif (N-X-S/T) required for N-linked glycosylation was aimed to insert into the sequence of the molecule at different positions, thus designing new analogous sequences of hEPO. For this purpose, the primary sequence and the three-dimensional structure of hEPO were studied thoroughly to select the compatible region for modification. The residues involved in receptor binding and those important for structural stability or bioactivity of hEPO were identified from previous literatures in order to keep them out of modification. Then every three amino acid stretch of the selected region was converted to the consensus tripeptide (N-X-S/T) by amino acid substitution at positions 0 and +2 of that stretch. Both N-X-S and N-X-T tripeptides were inserted at each position. In this way, multiple analogous sequences were created each of which was containing an additional N-glycosylation site along with the three native glycosylation sites.

### Prediction of glycosylation

The probability of glycosylation at both the modified site and the native sites of the new hEPO analogs was inspected by NetNGlyc 1.0 server (http://www.cbs.dtu.dk/services/NetNGlyc/) (Gupta & Brunak, 2002). For that, the FASTA sequences of the analogs were provided as input in the server. As output, the server detected and marked all the possible sites for addition of N-glycan moieties in the provided sequences and plotted their potential score against residue number in a graph. The potential score is the average output of nine neural networks and it ranges from 0 to 1. The neural networks are trained to differentiate between glycosylated and non-glycosylated asparagines by inspecting the surrounding sequence context of them. By default, the threshold for the potential score was set to 0.5 (50% of the total score). The analogous sequences which obtained >50% score for the prospective N-glycosylation sites were then used for 3D structure construction.

### Construction of 3D models of hEPO analogs

The three-dimensional (3D) models of the hEPO analogs were constructed by the homology modeling program named MODELLER version 9.18 (https://salilab.org/modeller/) (Eswar et al., 2008; Fiser & Šali, 2003; Šali & Blundell, 1993). The program implements the comparative protein structure modeling by satisfaction of spatial restraints. The input to run the program includes the amino acid sequence of the target molecule, the atomic coordinates of the templates, the target-template alignment and the script files containing necessary instructions. For the present study, the templates required for homology modeling were searched with the help of BLAST (Basic Local Alignment Search Tool) (https://blast.ncbi.nlm.nih.gov/Blast.cgi) (Altschul et al., 1990, 1997) against the Protein Data Bank database. To increase model accuracy, multiple templates were selected from the BLAST search. The PDB IDs of the selected templates were 1BUY, 1EER and 1CN4. A multiple sequence alignment was performed for these templates by MODELLER. Then, the target sequence of each hEPO analog was aligned to the sequences of the templates. Once the target-template alignment was constructed, MODELLER was run to build the 3D models of the hEPO analogs. The program was assigned to give five models for the target sequence along with their GA341 scores and DOPE (Discrete Optimized Protein Energy) scores. The GA341 score ranges from 0 to 1, where 0 represents models that tend to have an incorrect fold and 1 indicates models with proper folding which is comparable to X-ray structures. On the other hand, the DOPE score represents an atomic distance-dependent statistical potential used for initial assessment of the protein structure (Shen & Sali, 2006). Based on these initial assessment scores, the best model out of five models of each hEPO analog was selected. The constructed 3D models were then visualized by PyMOL (https://sourceforge.net/projects/pymol/) and UCSF Chimera (http://www.rbvi.ucsf.edu/chimera) (Pettersen et al., 2004) and subjected to further validation and analysis.

### Validation of model accuracy

The accuracy of the 3D models of the hEPO analogs generated by homology modeling was evaluated by a number of webservers. The stereochemical quality of the analogs was assessed by analyzing the Ramachandran plot generated by PROCHECK (Laskowski et al., 1993). When the atomic coordinates of each analog were provided to this server, the residue-by-residue geometry of the analog was checked and the phi/psi torsion angles of all residues were plotted in the Ramachandran plot (Ramachandran et al., 1963). The plot showed the distribution of all residues in four regions which were the most favored regions, additional allowed regions, generously allowed regions and disallowed regions. To detect any potential error in the computational models of the analogs ProSA-web (https://prosa.services.came.sbg.ac.at/prosa.php) (Wiederstein & Sippl, 2007) was utilized. The program calculated the quality score for the input structures of the analogs and displayed that in a plot in the context of native protein structures existing in Protein Data Bank. ERRAT (http://servicesn.mbi.ucla.edu/ERRAT/) (Colovos & Yeates, 1993) was used to analyze the statistics of non-bonded interactions between different atom types of the analogs and calculate the overall quality factor of the models by comparing with highly refined structures deposited in the Protein Data Bank. For structural comparison, each analogous structure was superimposed with the wild-type hEPO structure and the RMSD (root mean square deviation) value was calculated with the help of Tm-align server (https://zhanglab.ccmb.med.umich.edu/TM-align/) (Zhang & Skolnick, 2005).

### Incorporation of carbohydrate chains with hEPO analogs

To generate glycosylated models of the mutant analogs, the GlyProt server (http://www.glycosciences.de/glyprot/) (Bohne-Lang & Von der Lieth, 2005) was used for incorporating N-glycan molecules to the protein backbone. The database associated with the GlyProt server contains >1000 N-glycan structures constructed by SWEET-II program (Bohne et al., 1998) and optimized by TINKER MM3 force field (http://dasher.wustl.edu/tinker/). To attach appropriate glycan molecules with the analogs, the composition of the carbohydrate chains associated with the native hEPO was studied from previous literatures and one particular composition, among diversified structures, was selected for the present study. The desired carbohydrate structure was searched and selected from the database of GlyProt. The GlyProt server was then used to check whether the N-glycosylation sites of the analogs, both native and additional, were spatially accessible for glycosylation. Next, the carbohydrate moiety selected from the database was added to all the accessible N-glycosylation sites of the analogs, thus modeling the glycosylated form of the analogs.

### Molecular docking of hyperglycosylated analogs with the receptors

The interaction of the glycoengineered hEPO analogs with the erythropoietin receptors (EPORs) was inspected by molecular docking study performed by ZDOCK version 3.0.2 (http://zdock.umassmed.edu/) (Chen & Weng, 2003; Mintseris et al., 2007; Pierce et al., 2011). Since one molecule of native human erythropoietin interacts with two erythropoietin receptors (EPORs) *in vivo*, the dimer of EPOR was aimed to use as receptor in the docking study. The EPOR dimer was collected from the template 1EER which is a trimer of one hEPO molecule and two EPOR molecules. Before docking, 1EER was processed by PyMOL to remove the hEPO molecule and the water molecules, keeping only the two chains of EPOR. Then the EPOR dimer was provided to ZDOCK as receptor and the target hEPO analog as ligand to perform docking.

As output, the software returned a number of docked complexes along with their docking scores. The best docked complex of each analog was selected based on the highest score. The interaction interface of the selected hEPO analogs and EPORs was visualized and analyzed by PyMOL. The number of hydrogen bonds formed between the molecules was counted and the amino acids of both the analogs and the receptors involved in the bond formation were identified.

### Molecular dynamics (MD) simulation

To investigate the impact of amino acid mutation required for N-glycosylation on the structure and dynamic behavior of selected hEPO analogs, molecular dynamics simulation was performed in two phases – (a) for 50 ns and (b) for 100 ns. At first, 50 ns MD simulation was executed with the help of WebGRO software (https://simlab.uams.edu/) which uses GROMACS program for simulation (H. Bekker et al., 1993). The homology modeled structures of the analogs were provided as the initial structures for simulation. GROMOS96 43a1 forcefield was applied to prepare the protein structures for simulation (Chiu et al., 2009). The structure of each analog was solvated with the Simple Point Charge (SPC) water model in a triclinic box with fourier grid size of 36 × 40 × 36 Å. The system was neutralized by the addition of appropriate number of Na and Cl ions provided as 0.15 M salt solution. Following neutralization, energy minimization was done by steepest descent integrator applying 5000 steps. After that, the system was equilibrated under the NVT and NPT ensemble maintaining the constant temperature and pressure at 300 K and 1.0 bar respectively. The final MD simulation was run for 50 ns using leap-frog integrator (Hockney et al., 1974). During the course of simulation, Root Mean Square Deviation (RMSD), Radius of gyration (Rgyr), Root Mean Square Fluctuation (RMSF), Solvent Accessible Surface Area (SASA) and intramolecular hydrogen bond formation were computed using various GROMACS tools namely gmx rms, gmx gyrate, gmx rmsf, gmx sasa and gmx hbond, respectively. For comparing with the wild type molecule, the native hEPO structure (ID: 1buy) collected from RCSB PDB was also simulated and the trajectory of all molecules produced by the simulation was thoroughly analyzed.

The best potential analog was further analyzed by 100 ns MD simulation performed by Desmond module with OPLS_2005 forcefield from the Schrodinger suite of the Maestro non-commercial version 2021.1(Banks et al., 2005; Schrödinger, 2021). In this case, the analog-receptors docked complex and the wild-type hEPO-EPORs complex (PDB ID-1EER) were subjected to simulation run in order to investigate the stability of the complexes and evaluate their interaction. Before simulation, protein preparation was carried out for the analog-receptors and hEPO-EPORs complexes by PyMol and Swiss-PdbViwer respectively. Initially during simulation, each complex was solvated with TIP3P water model in an orthorhombic box by applying periodic boundary conditions with the margin set up at 10 Å. In order to maintain the physiological conditions, 0.15M NaCl salt was added to the solvated system to provide counter ions for neutralizing the total charge of the system. Then the system was run for 100 ps energy minimization through 5000 steps followed by equilibration. Finally, 100 ns MD simulation was implemented at a constant temperature of 310 K and a constant pressure of 1.013 bar by the NPT ensemble method. Various analyzing properties such as RMSD, RMSF, R_gyr_ and H-bonds were computed from the simulation trajectory to get critical insight about the complexes.

## Results

### Identifying suitable region to design new hEPO analogs

Human erythropoietin consists of four long alpha helices namely αA (residues 8-26), αB (residues 55-83), αC (residues 90-112), and αD (residues 138-161), two short helices namely αB’ (residues 47-52) and αC’ (residues 114-121), and two long and one short loops namely AB, CD and BC, respectively (Syed et al., 1998). According to previous literatures, the amino acids of hEPO which participate in high affinity and low affinity receptor binding, are resided in αA, αB’, αC and αD helices and AB loop of the molecule (Syed et al., 1998). No interacting residue is located in αB helix of hEPO. Therefore, αB helix was primarily selected as the suitable region for glycosylation in this study so that the modification in this region would not hamper the interaction of the molecule with the receptors. Among twenty-nine residues of αB helix, one residue (N^83^) is involved in natural N-glycosylation and three residues (E^62^, L^67^ and L^70^) are reported to affect folding and/or bioactivity of the molecule (Egrie et al., 1986; Elliott et al., 1997). So, these residues were not altered during modification of the sequence. All the other 25 residues of αB helix were considered to be transformed into N-X-S/T motif and 40 analogs were designed in this way which are listed in Table 1. Each mutated analog was carrying one extra N-glycosylation site in comparison to wild-type hEPO which holds maximum three N-linked carbohydrate chains.

**Table 1:** Potential analogs of hEPO with one additional N-glycosylation site. The probability of glycosylation at newly inserted asparagine (N) residues was estimated by NetNGlyc. Spatial accessibility of the analogs for glycosylation was investigated by GlyProt. Moleculer docking was performed by ZDOCK server. Hydrogen bonds of ten highest docking score obtaining analogs were counted and analysed by PyMOL. The number of hydrogen bonds formed by the analogs with the B chain and C chain of the receptors are listed here (shown in bracket) along with the total number of bonds. The best potential analogs selected for MD simulation are marked by bold.

| Analog | Amino acid changes | Probability of glycosylation (%) | Spatial accessibility | Docking score | Hydrogen bonds |
| --- | --- | --- | --- | --- | --- |
| Analog-55.1 | E55N-G57S | 81% | Yes | 1849.938 | - |
| Analog-55.2 | E55N-G57T | 83% | Yes | 1526.234 | - |
| Analog-56.1 | V56N-Q58S | 66% | Yes | 2313.094 | 15 (11, 4) |
| Analog-56.2 | V56N-Q58T | 72% | Yes | 2269.594 | 8 (7, 1) |
| Analog-57.1 | G57N-Q59S | 61% | Yes | 1631.982 | - |
| Analog-57.2 | G57N-Q59T | 68% | Yes | 990.955 | - |
| Analog-58.1 | Q58N-A60S | 62% | Yes | 1442.064 | - |
| Analog-58.2 | Q58N-A60T | 68% | Yes | 2111.764 | 6 (4, 2) |
| Analog-59.1 | Q59N-V61S | 58% | No | - | - |
| Analog-59.2 | Q59N-V61T | 65% | No | - | - |
| Analog-61.1 | V61N-V63S | 48% | - | - | - |
| Analog-61.2 | V61N-V63T | 55% | No | - | - |
| Analog-63.1 | V63N-Q65S | 36% | - | - | - |
| Analog-63.2 | V63N-Q65T | 43% | - | - | - |
| Analog-64.1 | W64N-G66S | 59% | Yes | Distorted structure | - |
| **Analog-64.2** | **W64N-G66T** | **66%** | **Yes** | **2050.646** | **20 (8, 12)** |
| Analog-66.1 | G66N-A68S | 58% | No | - | - |
| Analog-66.2 | G66N-A68T | 65% | No | - | - |
| Analog-69.1 | L69N | 70% | Yes | 2073.992 | 15 (11, 4) |
| Analog-69.2 | L69N-S71T | 75% | Yes | 1896.128 | - |
| **Analog-71.1** | **S71N-A73S** | **63%** | **Yes** | **2313.711** | **25 (13, 12)** |
| Analog-71.2 | S71N-A73T | 69% | Yes | 1914.648 | - |
| **Analog-72.1** | **E72N-V74S** | **65%** | **Yes** | **2051.840** | **19 (13, 6)** |
| Analog-72.2 | E72N-V74T | 71% | Yes | 2338.113 | 15 (11, 4) |
| Analog-73.1 | A73N-L75S | 65% | No | - | - |
| Analog-73.2 | A73N-L75T | 70% | No | - | - |
| Analog-74.1 | V74N-R76S | 64% | No | - | - |
| Analog-74.2 | V74N-R76T | 70% | No | - | - |
| Analog-75.1 | L75N-G77S | 60% | Yes | 1084.996 | - |
| Analog-75.2 | L75N-G77T | 66% | Yes | 1598.843 | - |
| **Analog-76.1** | **R76N-Q78S** | **52%** | **Yes** | **2025.016** | **24 (9, 15)** |
| Analog-76.2 | R76N-Q78T | 59% | Yes | 1251.386 | - |
| Analog-77.1 | G77N-A79S | 51% | No | - | - |
| Analog-77.2 | G77N-A79T | 58% | No | - | - |
| Analog-78.1 | Q78N-L80S | 66% | Yes | Distorted structure | - |
| Analog-78.2 | Q78N-L80T | 71% | Yes | Distorted structure | - |
| **Analog-79.1** | **A79N-L81S** | **63%** | **Yes** | **2068.152** | **20 (12, 8)** |
| Analog-79.2 | A79N-L81T | 69% | Yes | Distorted structure | - |
| Analog-80.1 | L80N-V82S | 61% | No | - | - |
| Analog-80.2 | L80N-V82T | 67% | No | - | - |

### Investigating probability of glycosylation in hEPO analogs

According to the prediction results obtained from NetNGlyc, thirty-seven of forty analogs obtained >50% potential score for N-glycosylation (Table 1). Three analogs namely Analog-61.1, Analog-63.1 and Analog-63.2 got <50% score, which were excluded from further analysis.

The glycosylation score of all thirty-seven analogs was found to be positive for both the native glycosylation sites and the newly introduced glycosylation sites. An example of glycosylation prediction for Analog-71.1 is shown in Supplementary Figure 1. The NetNGlyc server successfully detected four asparagine (N) containing N-X-S/T tripeptides at positions 24, 38, 71 and 83 in Analog-71.1 and represented their probability for carbohydrate addition by giving potential score for each site. Here, N^24^, N^38^ and N^83^ are native asparagines where glycosylation occurs naturally. Along with them, the newly introduced asparagine (N^71^) also attained potential score above the threshold level (0.6269), which indicated a possibility of an additional N-glycan molecule to be attached to this analog at position 71.

### Building 3D models of hEPO analog**s**

The modified sequences of thirty-seven analogs were used to build up the three-dimensional (3D) models through homology modeling. There was high sequence identity (≥95%) and 100% query coverage between the sequences of the templates and the analogs as found in the BLAST search. Most of the regions of these sequences were conserved as evident in the multiple sequence alignment, an example of which is exhibited for Analog-71.1 in Supplementary Figure 2. Based on the target-template alignment, five models of each analog were generated, all of which had same GA341 score i.e., 1.0, that represented native-like structure. However, The DOPE scores of the models were variable. The model that obtained the lowest value of the DOPE scores was selected as the best model. All the constructed models of the analogs were consisting of four long helices labeled as αA, αB, αC and αD, two short helical segments labeled as αB’ and αC’, and one short and two long loops. The loops connecting the helices were named as AB, BC and CD loops. An illustration of 3D model of Analog-71.1 is displayed in Figure 2A and its DOPE score and GA341 score are presented in Table 2.

**Figure 2:**
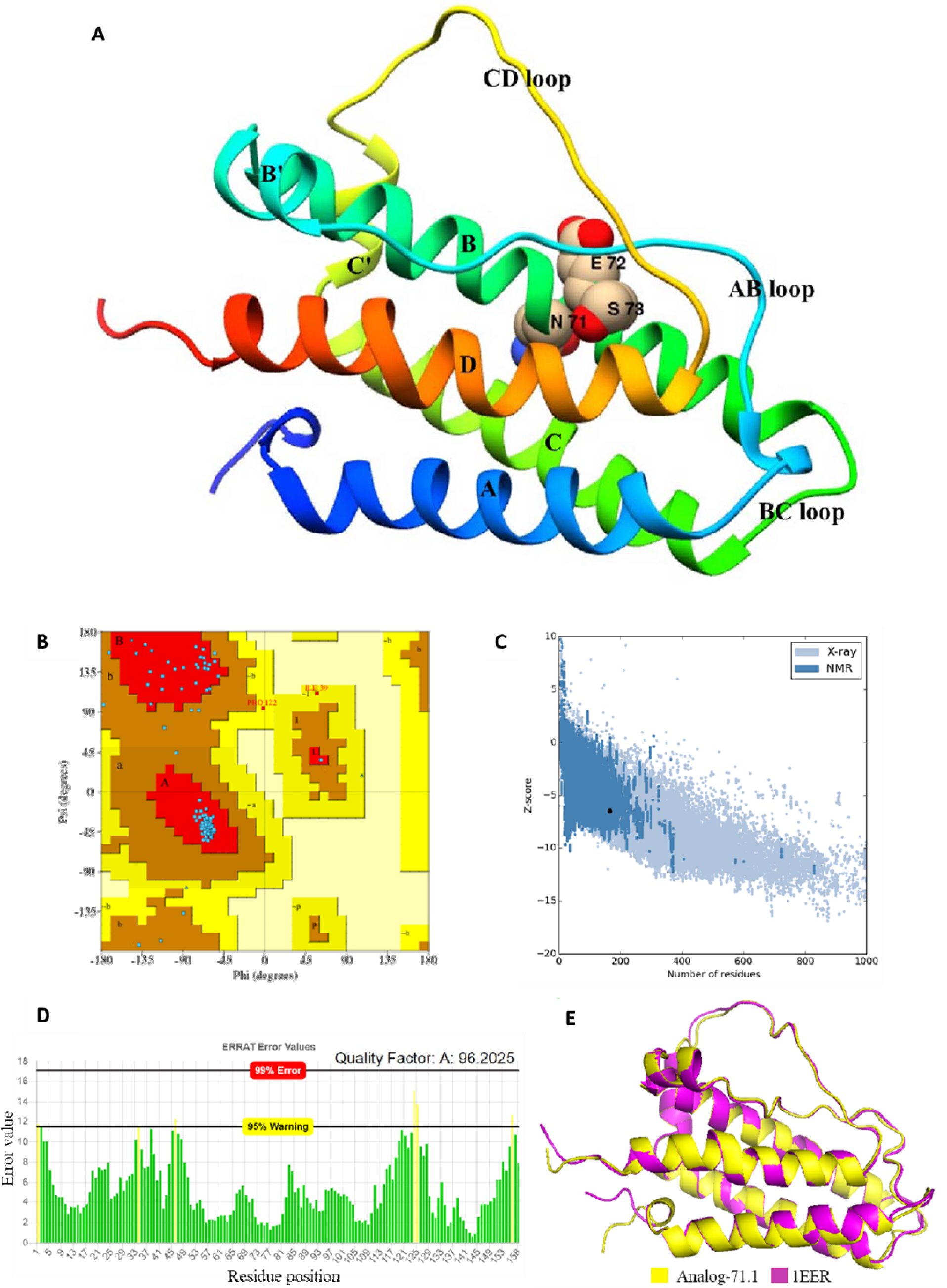
A) Three-dimensional model of Analog-71.1 generated by MODELLER through homology modeling and visualized by UCSF Chimera. The analog is represented by ribbon diagram and the modified region of the analog containing a probable glycosylation site (N71E72S73) is shown by sphere representation. Evaluation of 3D model of Analog-71.1. (B) Ramachandran plot obtained from PROCHECK showed that the distribution of maximum residues of the analog was in the most favored region. (C) The ProSA-web plot demonstrated that the Z-score of Analog-71.1 were justifiable and similar to the same sized protein structures of Protein Data Bank. (D) The overall quality factor of the analog calculated by ERRAT was very high and the error value of all the residues was within acceptable level. (E) The superimposition of the analog’s structure upon the template (1EER) structure performed by Tm-align server demonstrated that the two structures were highly similar with very little deviation.

**Table 2:** Evaluation scores of five models of Analog-71.1 generated by MODELLER. The model containing lowest DOPE score, indicated by bold, was selected as the best model of the analog.

| Models | DOPE score | GA341 score |
| --- | --- | --- |
| Analog-71.1.B99990001 | -18582.95898 | 1.00 |
| Analog-71.1.B99990002 | -18695.30469 | 1.00 |
| Analog-71.1.B99990003 | -18615.03320 | 1.00 |
| **Analog-71.1.B99990004** | **-18766.99609** | **1.00** |
| Analog-71.1.B99990005 | -18383.65039 | 1.00 |

### Evaluation of the 3D models

The evaluation of the 3D models of the hEPO analogs was performed by various *in silico* tools, the results of which are summarized and presented in Supplementary Table 1. Ramachandran plot was generated for the modeled analogs by the program PROCHECK to analyze the phi/psi torsion angles of all the residues of those models. Most of the analogs had >90% residues in the most favored regions. Majority of them had no residue in the disallowed regions, while others had only 0.7% residues in that region. The inclusive analysis of the Ramachandran plot confirmed good geometry of the 3D models of the analogs. According to the results obtained from ProSA-web, the Z-scores of the analogs were between −5.92 and −6.75, which were resided within the acceptable range of scores of the experimental protein structures. The overall quality factor of the analogs was then calculated by ERRAT. The ERRAT score of all the analogs was acceptable, except one analog i.e., Analog-76.2, the quality factor of which was <80. Two analogs namely Analog-69.1 and Analog-71.1 secured the most satisfying quality factor which was >95. All the residues of these analogs had error values below 99% rejection limit. Lastly the homology modeled structure of each analog was compared with the native hEPO structure by superimposition to investigate the deviation of the analogs. The RMSD (root mean square deviation) value of all the analogs was below 1.5 nm. The little deviation of the analogs ensured their high similarity with the natural structure. As an example, the evaluation parameters for Analog-71.1 are presented in Figure 2 (B-E) and Table 3. The Ramachandran plot analysis showed that Analog-71.1 had 92.5%, 6.8% and 0.7% residues in the most favored, additional allowed and generously allowed regions, respectively and no amino acid in the disallowed regions. The ProSA-web Z-score of the analog was −6.48 which belonged to the allowable range. The ERRAT quality score of the analog was very high (96.2025) and the plot demonstrated no region with significant error. The superimposed structure of Analo-71.1 with the wild-type structure (1EER) had very little RMSD (1.27).

**Table 3:** Evaluation parameters of homology model of Analog-71.1.

| Analog | Ramachandran plot |  |  |  | Z-score | Quality factor | Whole RMSD (nm) |
| --- | --- | --- | --- | --- | --- | --- | --- |
|  | Most favored regions | Additional allowed regions | Generously allowed regions | Disallowed regions |  |  |  |
| Analog-71.1 | 92.5% | 6.8% | 0.7% | 0.0% | -6.48 | 96.2025 | 1.27 |

### Constructing glycosylated models of the hEPO analogs

After homology modeling and model validation, the analogs were examined by GlyProt to check the spatial accessibility of their newly introduced N-glycosylation sites in 3D configuration. The glycosylation sites were found to be accessible in twenty-four analogs among thirty-seven analogs. The glycan moiety to be attached at these sites was a four branched carbohydrate chain chosen from previous literature (Sasaki et al., 1987) and selected from the database of GlyProt (Figure 3A). It is composed of 17 monosaccharides comprising one fucose, four galactoses, three mannoses, six N-acetyleglucoses and three N-acetyleneuraminic acids (sialic acids). The first three branches of this tetraantennary glycan have sialic acids as the terminal molecules, whereas the fourth branch is ended with galactose. Each sialic acid is linked to the branch by 2→3 glycosidic linkage. The GlyProt server constructed the glycosylated models of the analogs by incorporating this N-linked carbohydrate chain both at the native glycosylation sites and the additionally inserted glycosylation sites of the analogs. An illustration of hyperglycosylated form of Analog-71.1 is demonstrated in Figure 3B which had four glycans at N^24^, N^38^, N^71^ and N^83^.

**Figure 3:**
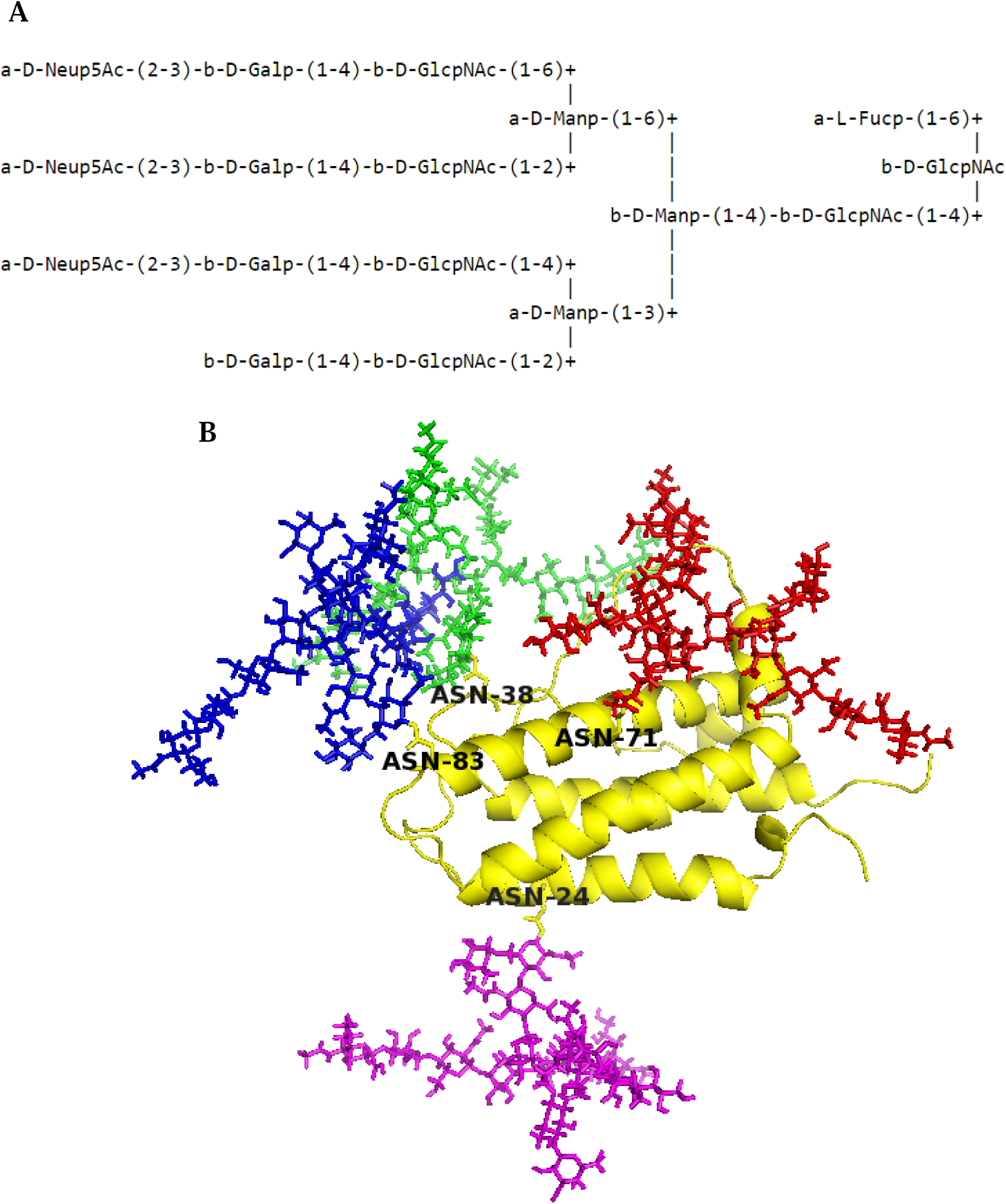
Modeling of glycosylated form of Analog-71.1. (A) The tetraantennary carbohydrate chain selected for glycoengineering from GlyProt server containing three sialic acids as the terminal molecules of three branches. (B) Hyperglycosylated structure of Analog-71.1 containing four carbohydrate chains at the native and additional glycosylation sites. The carbohydrate chain attached to the asparagine (N^71^) of the additional glycosylation site is represented by red color, whereas the native glycans are represented by purple (N^24^), green (N^38^) and blue (N^83^) colors.

### Interaction analysis of glycosylated hEPO analogs with the receptors

The hyperglycosylated models of twenty-four analogs were docked with erythropoietin receptors (EPORs) with the help of ZDOCK to investigate their interaction. Four out of twenty-four analogs formed unusual docked complexes (Table 1). This indicated that their sequence modification and position of carbohydrate addition made them unfit to interact with the receptors. So, these analogs were disqualified for next analysis. Each of the remaining twenty analogs got attached with the EPOR dimer and formed an appropriate tri-body complex structure like the native hormone-receptor complex. For instance, the docked complex of Analog-71.1 and the receptors is displayed in Figure 4. The analogs were then scrutinized on the basis of their docking score and bond formation in order to evaluate how strongly and properly they have bound to the receptors.

**Figure 4:**
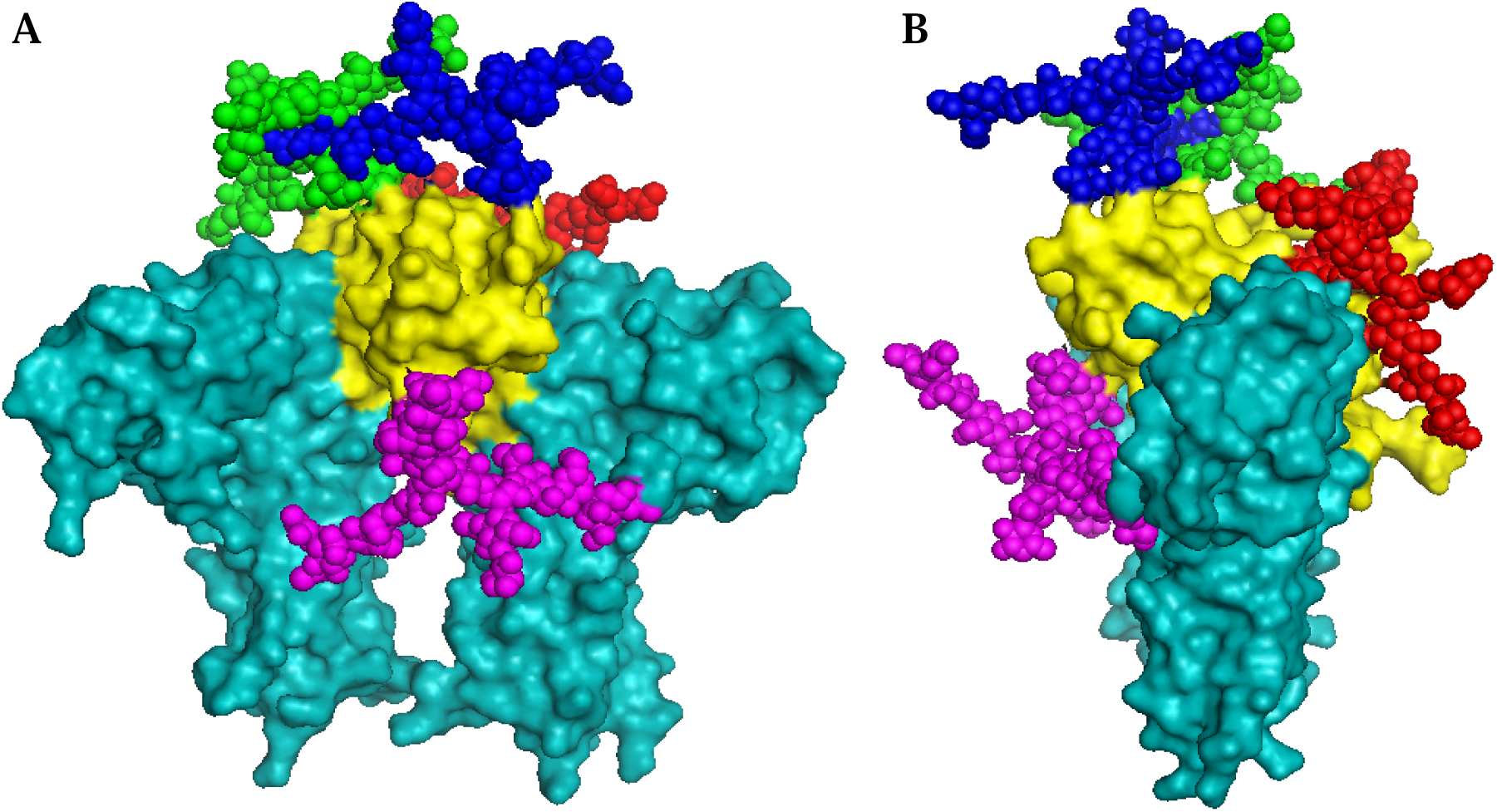
Docked complex of glycosylated Analog-71.1 and EPORs dimer; A (front view) and B (side view) showing surface and sphere representation of the complex. Analog-71.1 and the receptors are presented by yellow and cyan colors, respectively. The carbohydrate chains are presented by red (N^71^), purple (N^24^), green (N^38^) and blue (N^83^) colors.

Among the twenty analogs, some obtained very high docking score such as Analog-72.2 (2338.113), some had moderate score such as Analog-55.2 (1526.234), while some got very low score such as Analog-57.2 (990.955) (Table 1). According to the ZDOCK score, ten highest score obtaining analogs were selected to study their interaction interface with the receptors more precisely. The number of hydrogen bonds formed within these docked complexes is listed in Table 1. The study revealed that two analogs had lowest number of hydrogen bonds with the receptors which were Analog-56.2 (8) and Analog-58.2 (6). Three analogs formed moderate number of hydrogen bonds with the receptors which were Analog-69.1 (15), Analog-56.1 (15) and Analog-72.2 (15). These analogs were considered as less potential and excluded from next experiment. Finally, five analogs, with higher number of hydrogen bonds along with higher docking score, were qualified for molecular dynamics simulation study. The qualified analogs were Analog-64.2, Analog-71.1, Analog-72.1, Analog-76.1 and Analog-79.1 and the quantity of hydrogen bonds occurring within their docked complexes were 20, 25, 19, 24 and 20, respectively. Evidently, Analog-71.1 had the highest amount of bonds formed with the receptors. The interaction interface of this analog is demonstrated in Figure 5 and discussed in details below.

**Figure 5:**
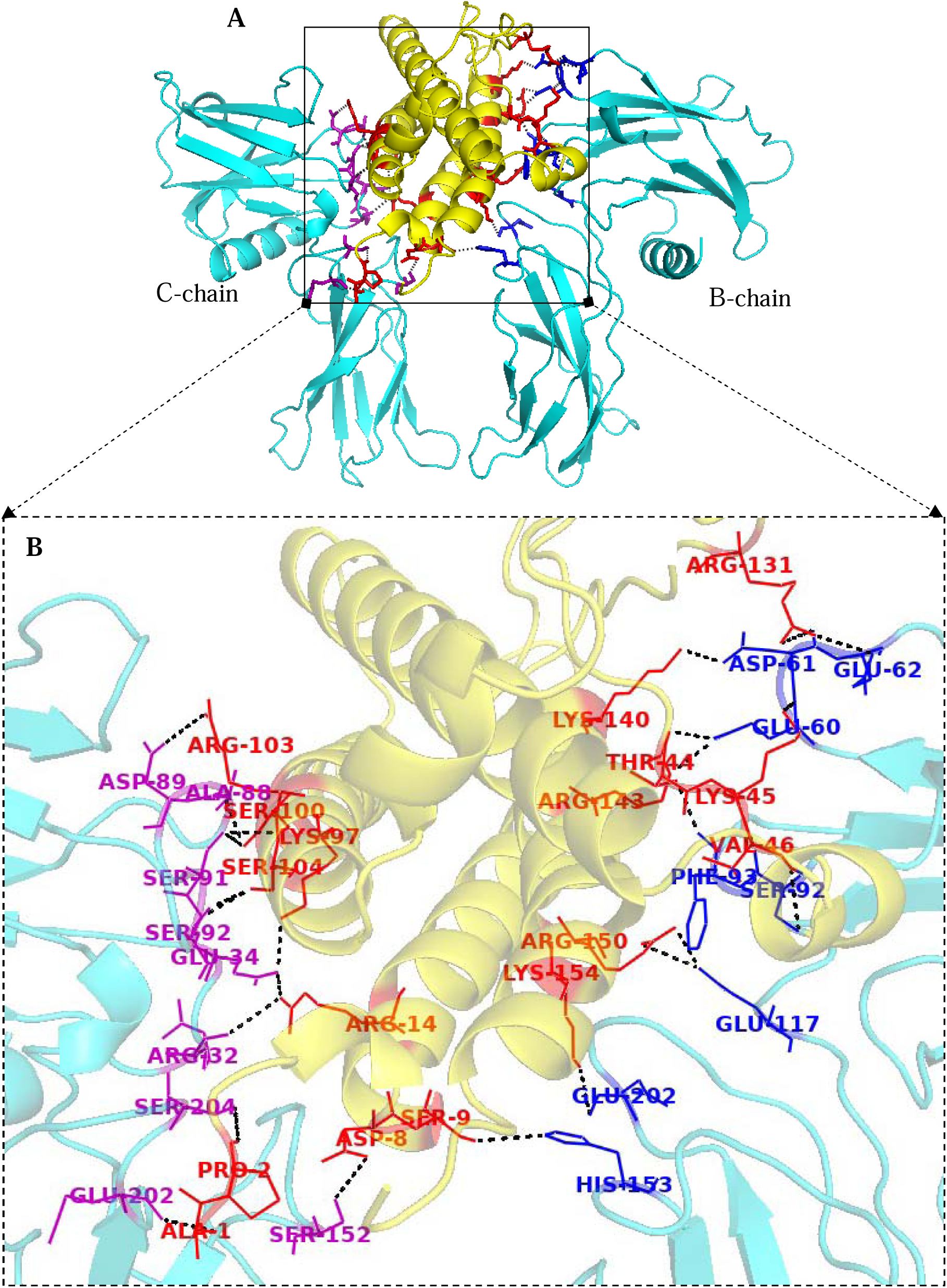
Interaction interface between Analog-71.1 and EPORs. (A) Ribbon diagram showing Analog-71.1-EPORs docked complex where interacting residues are displayed by stick view. (B) Enlarged view of interaction interface; red-, blue- and purple-colored lines representing the amino acids of Analog-71.1, EPOR B-chain and EPOR C-chain, respectively; black dots representing the hydrogen bonds formed between the amino acids.

The ZDOCK score acquired by the docked complex of Analog-71.1 and EPORs was 2313.711 which was second highest among all the analogs. The comprehensive examination of its interaction interface discovered that 13 hydrogen bonds, out of 25, were formed between the analog and the B-chain of the receptor forming higher affinity interface, while 12 hydrogen bonds were formed between the analog and the C-chain of the receptor forming lower affinity interface. For comparative analysis, the hydrogen bonds formed within the native hEPO-EPORs complex structure (1EER) were also analyzed which had 29 bonds between hEPO and receptor molecules (Supplementary Figure 3). Here, the wild-type hEPO had 16 hydrogen bonds with the B-chain and 13 with the C-chain of the receptors. The amino acid residues involved in hydrogen bond formation in both analog-receptor complex and native hEPO-receptor complex are indexed in Table 4.

**Table 4:**
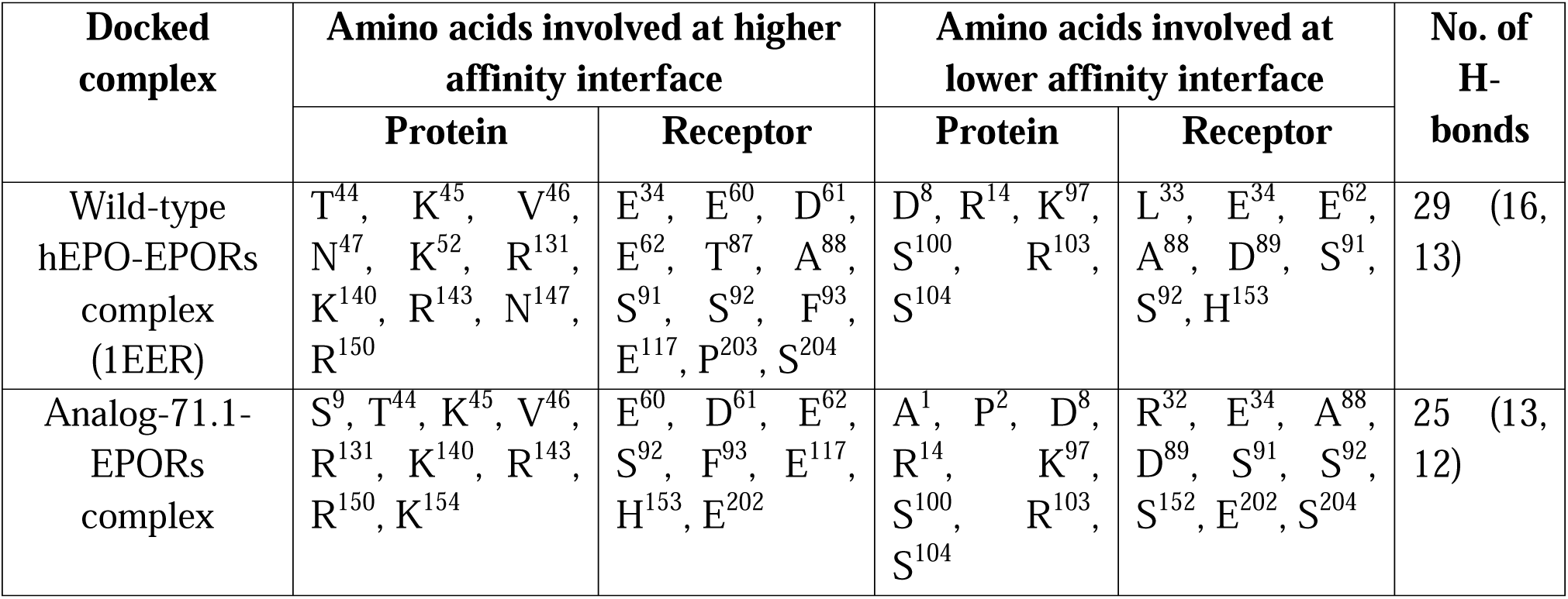
Amino acids resided at the interaction interface of the docked complexes and number of hydrogen bonds formed among them.

| Docked complex | Amino acids involved at higher affinity interface |  | Amino acids involved at lower affinity interface |  | No. of H-bonds |
| --- | --- | --- | --- | --- | --- |
|  | Protein | Receptor | Protein | Receptor |  |
| Wild-type hEPO-EPORs complex (1EER) | T<sup>44</sup>, K<sup>45</sup>, V<sup>46</sup>, N<sup>47</sup>, K<sup>52</sup>, R<sup>131</sup>, K<sup>140</sup>, R<sup>143</sup>, N<sup>147</sup>, R<sup>150</sup> | E<sup>34</sup>, E<sup>60</sup>, D<sup>61</sup>, E<sup>62</sup>, T<sup>87</sup>, A<sup>88</sup>, S<sup>91</sup>, S<sup>92</sup>, F<sup>93</sup>, E<sup>117</sup>, P<sup>203</sup>, S<sup>204</sup> | D<sup>8</sup>, R<sup>14</sup>, K<sup>97</sup>, S<sup>100</sup>, R<sup>103</sup>, S<sup>104</sup> | L<sup>33</sup>, E<sup>34</sup>, E<sup>62</sup>, A<sup>88</sup>, D<sup>89</sup>, S<sup>91</sup>, S<sup>92</sup>, H<sup>153</sup> | 29 (16, 13) |
| Analog-71.1-EPORs complex | S<sup>9</sup>, T<sup>44</sup>, K<sup>45</sup>, V<sup>46</sup>, R<sup>131</sup>, K<sup>140</sup>, R<sup>143</sup>, R<sup>150</sup>, K<sup>154</sup> | E<sup>60</sup>, D<sup>61</sup>, E<sup>62</sup>, S<sup>92</sup>, F<sup>93</sup>, E<sup>117</sup>, H<sup>153</sup>, E<sup>202</sup> | A<sup>1</sup>, P<sup>2</sup>, D<sup>8</sup>, R<sup>14</sup>, K<sup>97</sup>, S<sup>100</sup>, R<sup>103</sup>, S<sup>104</sup> | R<sup>32</sup>, E<sup>34</sup>, A<sup>88</sup>, D<sup>89</sup>, S<sup>91</sup>, S<sup>92</sup>, S<sup>152</sup>, E<sup>202</sup>, S<sup>204</sup> | 25 (13, 12) |

The amino acids of Analog-71.1 present at higher affinity interface were S^9^, T^44^, K^45^, V^46^, R^131^, K^140^, R^143^, R^150^ and K^154^. On the other hand, the amino acids of B-chain of EPORs present at higher affinity interface were E^60^, D^61^, E^62^, S^92^, F^93^, E^117^, H^153^ and E^202^. Among these residues, S^9^ interacting with H^153^ and K^154^ interacting with E^202^ were not present in the interaction interface of wild-type hEPO-EPORs. Apart from these four residues, the rest of analog-receptors complex were common with that of wt-hEPO-EPORs. The amino acids of Analog-71.1 present at lower affinity interface were A^1^, P^2^, D^8^, R^14^, K^97^, S^100^, R^103^ and S^104^ and the amino acids of EPORs interacting with those residues were R^32^, E^34^, A^88^, D^89^, S^91^, S^92^, S^152^, E^202^ and S^204^. Among these residues A^1^, P^2^ of the analog and R^32^, E^202^ of EPOR C-chain were not present in the interaction interface of wild-type complex. Apart from these four residues, the rest were all common with wild-type complex. This study indicated that the interaction pattern of Analog-71.1 with the receptors was highly similar to the native hEPO molecule. Moreover, the high docking score and highest bond formation manifested very strong interaction of the analog with the receptors.

### Molecular dynamics simulation study of selected hEPO analogs

According to molecular docking study and other analysis mentioned earlier, five analogs were selected for molecular dynamics simulation in order to inspect how the amino acid substitution for N-glycosylation would affect the structural stability of the analogs in a dynamic environment. The selected analogs were Analog-64.2, Analog-71.1, Analog-72.1, Analog-76.1 and Analog-79.1. The MD simulation was initially conducted for 50 ns by WebGRO software that provided trajectories of RMSD, RMSF, R_gyr_, SASA and hydrogen bonds of the analogs which were rationally assessed and compared with the respective trajectories of wild-type hEPO.

### Molecular stability analysis

RMSD (Root mean square deviation) is a reliable indicator of protein’s structural stability. In case of MD simulation, RMSD represents variation of backbone atoms of a protein from its initial structure to its final conformation (Aier et al., 2016). In the present study the RMSD of the wild-type hEPO and the mutated analogs were calculated and plotted against 50 ns timeframe which is presented in Figure 6 and summarized in Table 5. In the beginning, all simulated system including the wild-type hEPO showed a rapid increase in RMSD values. At about 10 ns the wild-type converged to equilibrium phase and maintained a constant RMSD around 0.381 ± 0.024 till the end of the simulation. Although it showed little increase around 35 ns, but then maintained the plateau. The equilibrium state with smaller deviation of the wild-type molecule indicates its stable conformation during the course of MD simulation.

**Figure 6:**
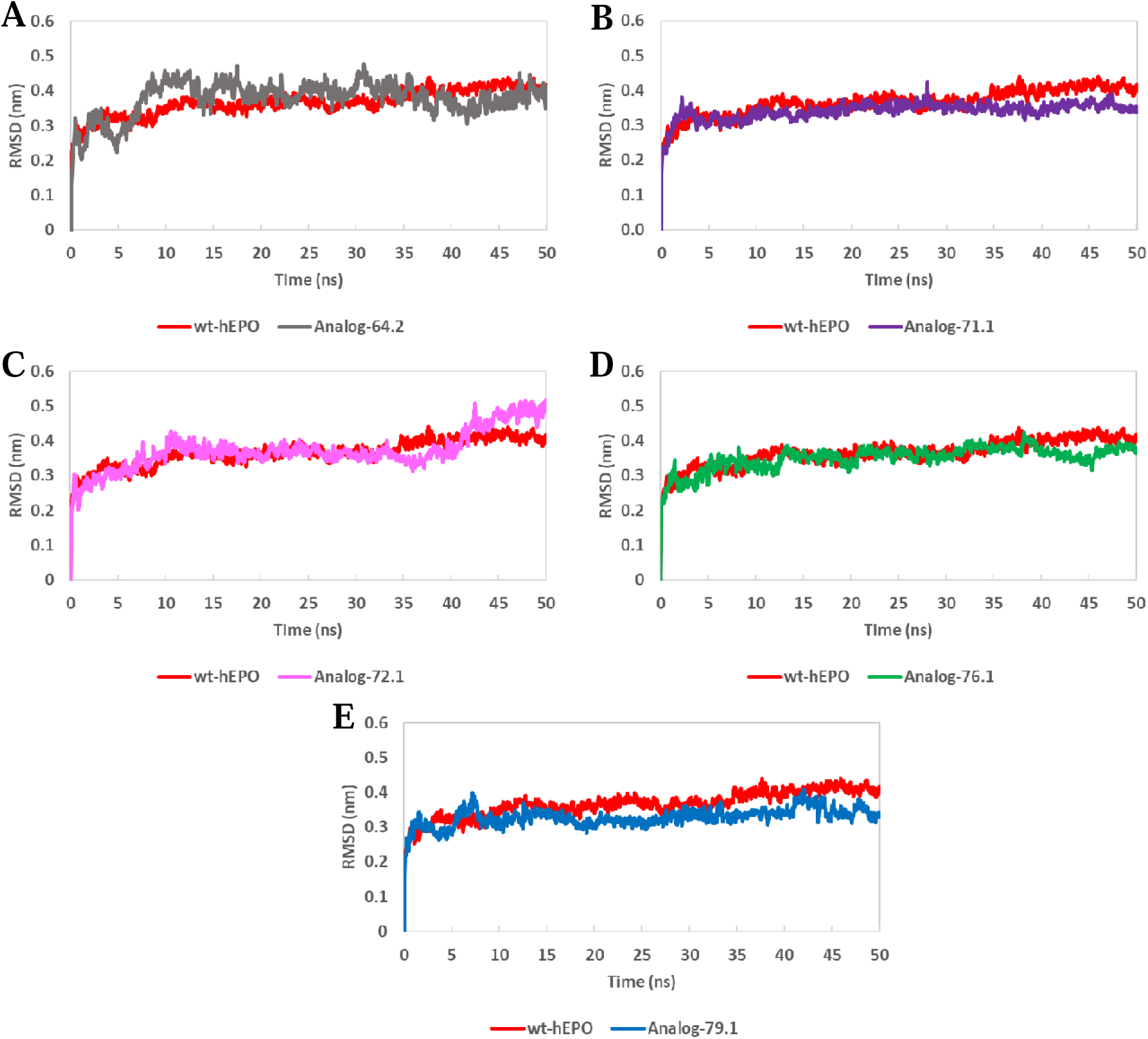
RMSD plots of Analog-64.2 (A), Analog-71.1 (B), Analog-72.1 (C), Analog-76.1 (D) and Analog-79.1 (E) in comparison to wild-type hEPO obtained from 50 ns MD simulation. Analog-71.1 and Analog-76.1 showed relatively more stable RMSD trajectory than other analogs.

**Table 5:** Molecular dynamics simulation study for wild-type hEPO and mutated analogs. The average value and standard deviation of RMSD, Rgyr, H-bonds and SASA of wild-type and the analogs were calculated over 50 ns simulation. The RMSF profiles of the molecules were analyzed to identify the highly fluctuating residues.

| Wild-type and analogs | Average RMSD over the equilibration phase | Highly fluctuating residues | Average $R_{\text{gyr}}$ over the equilibration phase | Average hydrogen bonds | Average SASA over the equilibration phase |
| --- | --- | --- | --- | --- | --- |
| Wt-hEPO | $0.381 \pm 0.024$ | Absent | $1.594 \pm 0.014$ | 119 | $89.842 \pm 2.272$ |
| Analog-64.2 | Unstable | Present | N/A | N/A | N/A |
| Analog-71.1 | $0.345 \pm 0.017$ | Absent | $1.629 \pm 0.008$ | 124 | $87.968 \pm 2.578$ |
| Analog-72.1 | Unstable | Present | N/A | N/A | N/A |
| Analog-76.1 | $0.366 \pm 0.017$ | Present | $1.598 \pm 0.013$ | 121 | $80.396 \pm 2.569$ |
| Analog-79.1 | Unstable | Present | N/A | N/A | N/A |

Among the simulated analogs, Analog-64.2 didn’t reach the equilibrium state at all and displayed large fluctuations throughout the simulation trajectory (Figure 6A). Analog-72.1 attained equilibration for some time during simulation ( 11-40 ns), but then it gradually followed an upward trend indicating higher structural deviation (Figure 6C). Analog-79.1 also achieved plateau for limited time of simulation ( 9-40 ns). But after 40 ns, higher fluctuations were observed in its RMSD values which speculated instability of the analog at the later phase of the simulation (Figure 6E).

The remaining two analogs, namely Analog-76.1 and Analog-71.1, demonstrated stable RMSD trajectory during simulation (Figure 6B and 6D). Initially the RMSD values of both analogs rose abruptly just like the native hEPO, which eventually became stabilized. After some initial fluctuation, Analog-76.1 reached plateau at 13 ns. The average RMSD of this analog was 0.359 ± 0.022. After 40 ns, the analog showed a gradual decrease followed by increase in RMSD values (Figure 6D). However, this fluctuation of Analog-76.1 was not as prominent as Analog-79.1. On the other hand, Anaog-71.1 attained equilibration at 5 ns, which remained constant around 0.345 ± 0.017 till the end of the simulation. During the equilibrium phase, there is only one point (28 ns) when the trajectory of the analog showed a highest peak of 0.424 nm, but after that the analog maintained the plateau. Particularly from 35 ns, the analog kept continuously lower RMSD values than the wild-type for the rest of the simulation, indicating comparatively less deviation and more stability than the wild-type molecule (Figure 6B).

### Residual flexibility analysis

RMSF profile of a protein indicates how much the residues of the protein fluctuate from their original mean positions during MD simulation(Bagewadi et al., 2023). More fluctuation implies more flexibility of the structure. Calculating RMSF of the analogs allowed us to detect the flexible regions of the analogs and compare that with the wild-type molecule. The study of RMSF profile of wild-type hEPO revealed that the helical regions of the molecule had relatively less fluctuation whereas the loop regions had more fluctuation (Figure 7). Also, the residues of N and C terminal of the molecule displayed higher flexibility.

**Figure 7:**
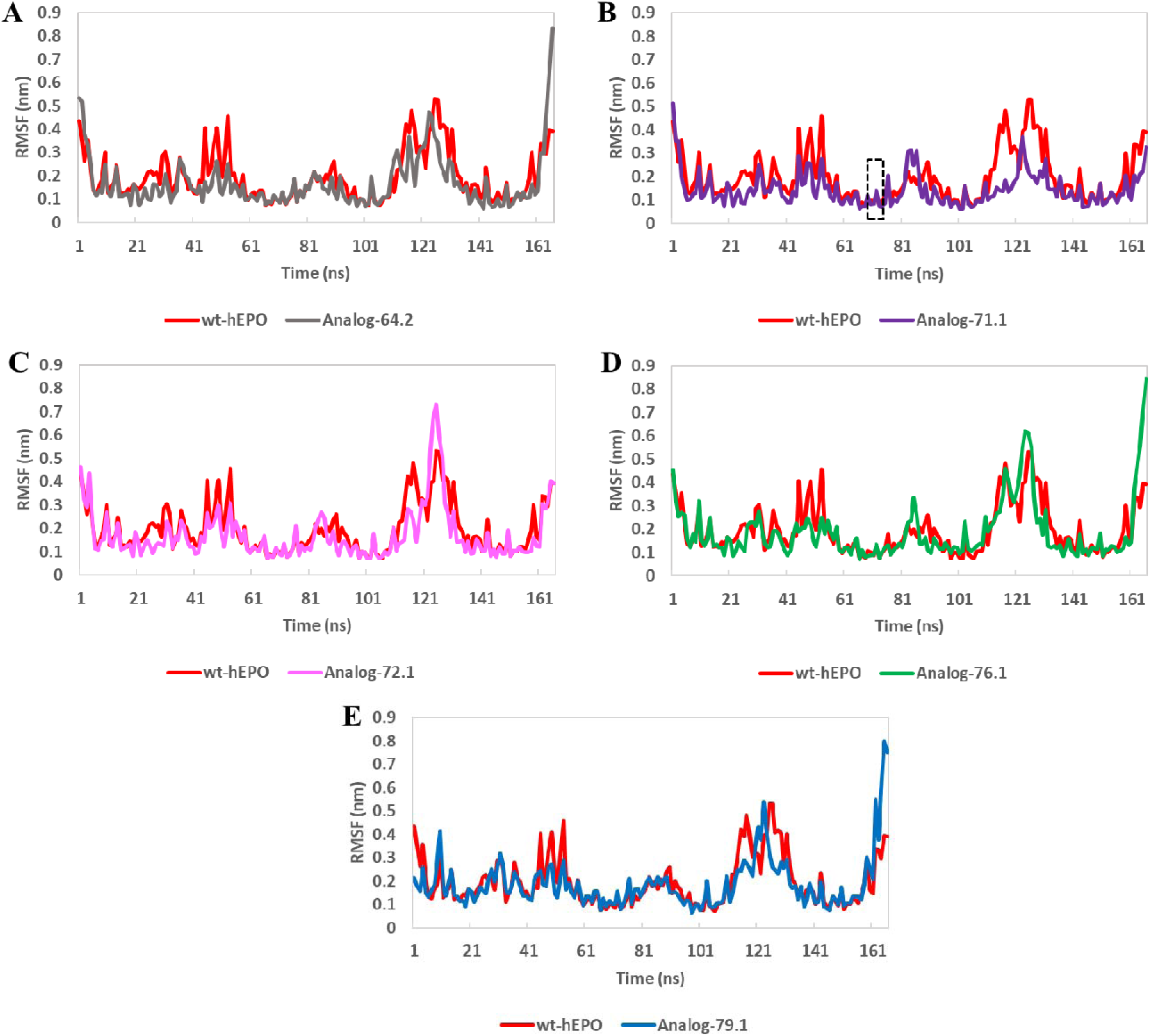
RMSF profiles of Analog-64.2 (A), Analog-71.1 (B), Analog-72.1 (C), Analog-76.1 (D) and Analog-79.1 (E) in comparison to wild-type hEPO produced by 50 ns MD simulation. All the analogs, except Analog-71.1, displayed some highly fluctuating residues relative to their native counterparts. Analog-71.1 showed relatively lower residual oscillation than wild-type indicating less flexibility of the structure. In addition, the modified region of Analog-71.1 (shown by black dotted box) didn’t show any significant variation than wt-hEPO.

The degree of residual fluctuation of Analog-71.1, which had very stable RMSD trajectory, was aimed to measure. Figure 7B demonstrates the alignment of RMSF profiles of Analog-71.1 and native hEPO. As the plot depicted, the values of analog’s RMSF were mostly lower than the native molecule, which indicated less flexibility of the analog structure and supported the result of RMSD. The mutated amino acids of the analog (S71N and A73S) were examined more precisely which revealed slightly lower fluctuation in the analog compared to wt-hEPO, articulating that the residual modification didn’t cause more flexibility, rather kept stability in the analog. The regions of the analog that showed higher variation relative to their native counterparts were the N-terminal residue and the residues of position 83-86 (Figure 7B). Here, the residue at position 83 (N^83^) is structurally important since it is a natural N-glycosylation site.

But when the wild-type hEPO structure was deposited in the Protein Data Bank through NMR spectroscopy method, asparagine (N^83^) at this position was replaced by lysine (K^83^) (Cheetham et al., 1998). So, variation at this position between the analog and the wild-type is expected. Moreover, the fluctuation of 83-86 residues was not very significant and the residues belong to loop region, so little fluctuation is acceptable.

Among the other mutant analogs, Analog-64.2, Analog-76.1 and Analog-79.1 had major fluctuation in their C terminal region which was much higher than the wild-type (Figure 7 (A, D, E)). Highly flexible region, relative to wild-type, was also observed in Analog-72.1 at position 123-126 (Figure 7C). Here, Ser^126^ is a structurally important amino acid as it is involved in O-glycosylation in native hEPO. Since Analog-64.2, Analog-72.1 and Analog-79.1 showed inconsistency in RMSD trajectory and also highly fluctuating region in RMSF profiles, these analogs were excluded from further analysis.

### Structural compactness analysis

The compactness of a protein structure can be measured by radius of gyration (R_gyr_), relatively constant values of which, during simulation, indicate that the protein has stably folded conformation (Ghasemi et al., 2016). The R_gyr_ analysis for the wild-type hEPO and the remaining stable analogs were performed in order to inspect whether the amino acid substitution would alter the structural compactness of the analogs. The R_gyr_ values of both wild-type and the mutants started with □1.68 nm followed by some major fluctuation in the beginning of the trajectory (Figure 8(A and B)). Around 2 ns the wt-hEPO was equilibrated maintaining an average R_gyr_ of 1.594 ± 0.014 nm till the end of the simulation. At 35 ns the wild-type R_gyr_ values were slightly decreased, but then continued with a plateau. On the other hand, equilibrium state was observed for both Analog-71.1 and Analog-76.1 from 5 ns to the termination of the trajectory, keeping the mean R_gyr_ at 1.629 ± 0.008 and 1.598 ± 0.013 respectively (Table 5). However, the trajectory of Analog-76.1 was not as steady as Analog-71.1. From 30 ns the R_gyr_ of Analog-76.1 was gradually reduced and then again elevated from 40 ns making a wavy trajectory (Figure 8B). Although Analog-71.1 maintained relatively higher R_gyr_ values than both wt-hEPO and Analog-76.1, the values of the analog were more stable with less fluctuations throughout the simulation (Figure 8A).

**Figure 8:**
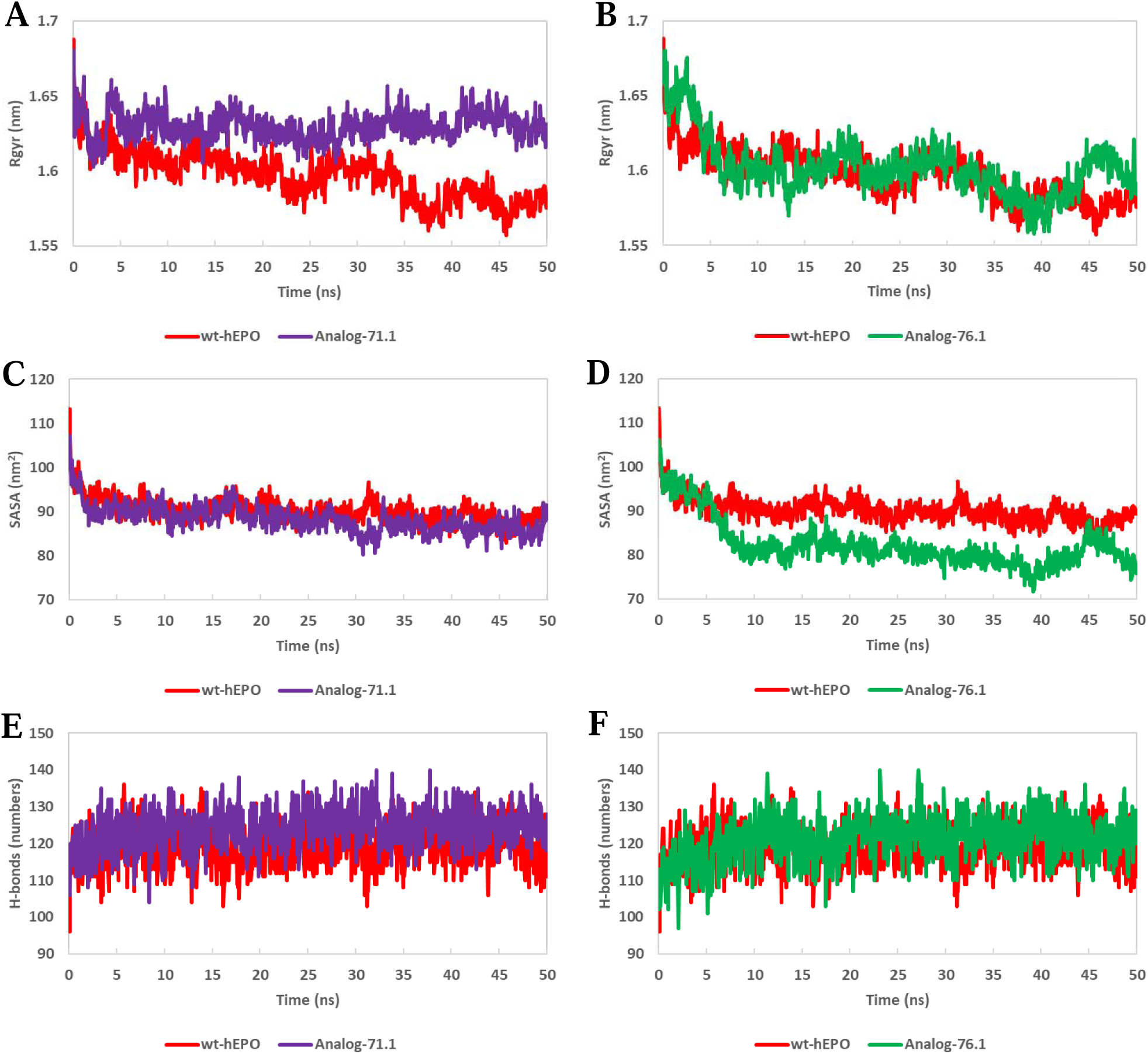
The values of R_gyr_ (A, B), SASA (C, D) and hydrogen bonds (E, F) plotted against simulation time for Analog-71.1 and Analog-76.1, respectively, in comparison to wt-hEPO. The analysis of the trajectories demonstrated that Analog-71.1 showed more stability than Analog-76.1 during 50 ns simulation.

### SASA analysis

Another determining factor of protein stability is SASA (solvent accessible surface area), which is defined by the exposed area of the protein responsible for making interactions with the surrounding solvent molecules (Borjian Boroujeni et al., 2021). As shown in Figure 8C, Analog-71.1 displayed a steady trajectory from 2 ns till the end of the simulation just like the wild-type, with little variation at 30-33 ns. The average SASA of the analog (87.968 ± 2.578) was also very close to the wild-type (89.842 ± 2.272) (Table 5). On the other hand, the trajectory of Analog-76.1 for SASA was not constant throughout the simulation. The analog achieved plateau at 9 ns which became relatively unstable after 38 ns (Figure 8D). This result was similar with the analysis of RMSD and R_gyr_ of the analog (discussed earlier). The average SASA of the analog (80.396 ± 2.569) also varied largely from the wt-hEPO (Table 5).

### Analysis of hydrogen bond formation

The number of intramolecular hydrogen bonds is crucial for stabilizing the conformation of a polypeptide (Chikalov et al., 2011). Therefore, hydrogen bonds of both native hEPO and the selected analogs were estimated which is presented against the simulation time in Figure 8(E and F) and summarized in Table 5. As the H-bond plot demonstrated, no significant variation was found between the analogs and the original hEPO.

### Molecular dynamics simulation study of analog-receptor docked complex

Among forty hEPO analogs designed in this study, five potential analogs were analyzed through 50 ns MD simulation, the findings of which manifested Analog-71.1 as the most stable analog after amino acid mutation. Further in-depth investigation on Analog-71.1 was aimed to conduct by extending the simulation time to 100 ns. In this experiment, the docked complex of Analog-71.1 with EPORs dimer was simulated in order to scrutinize the dynamic interaction of the analog with the receptors. Subsequently the conformational stability of the entire analog-receptor complex structure over time was unraveled through this investigation. Desmond simulation package from Schrodinger was utilized in this purpose. The necessary evaluation parameters like RMSD, R_gyr_, intramolecular hydrogen bonds formation and RMSF were calculated from the simulation trajectory which are presented in Figure 11 and 12 in comparison to native hEPO-ERORs complex.

The RMSD plot was generated from 100 ns simulation for the backbone atoms of both Analog-71.1-EPORs complex and wild-type hEPO-EPORs complex. According to the RMSD plot, the wt-hEPO-EPORs complex achieved plateau at 2 ns which took a leap after 37 ns and then the plateau was continued for the rest of the trajectory (Figure 9A). The average RMSD of the equilibrated wild-type complex was 3.444 ± 0.391 nm. On the other hand, Analog-71.1-EPORs docked complex exhibited major fluctuations at the beginning of the simulation until 10 ns. After that the system was equilibrated which showed a slight reduction at 48 ns followed by a little increase after 68 ns, none of which was much significant to point toward instability (Figure 9A). Rather the consistent RMSD trajectory of the analog-receptors complex with an average of 3.465 ± 0.239 nm highlighted the complex’s stability with fewer conformational changes.

**Figure 9:**
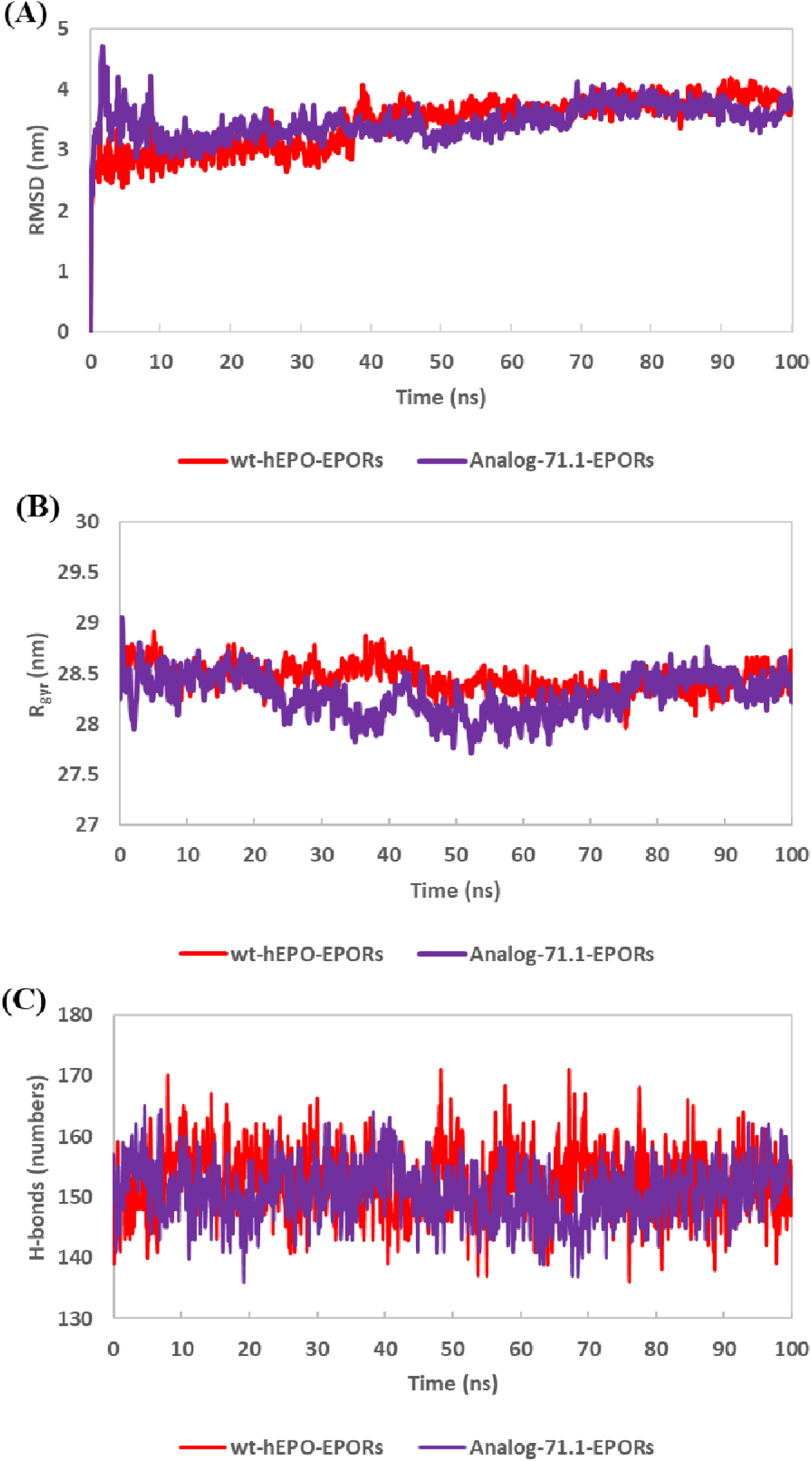
100 ns MD simulation trajectories displaying RMSD (A), R_gyr_ (B) and H-bonds (C) plots of Analog-71.1-EPORs complex (purple) along with wt-hEPO-EPORs complex (red).

To substantiate the structural stability of Analog-71.1-EPORs complex, further investigation was executed on its R_gyr_ profile which is illustrated along with the R_gyr_ of wild-type complex in Figure 9B. The wild-type hEPO-EPORs complex demonstrated a steady trajectory for R_gyr_ throughout the simulation keeping an average value of 28.451 ± 0.136 nm. For Analog-71.1-EPORs complex, the R_gyr_ trajectory started with major fluctuation followed by a plateau from 6 to 22 ns and from 55 to 75 ns. A major fluctuation pattern was observer with increase and reduction of R_gyr_ values within 23-74 ns which indicated structural modification. However, the analog-receptors complex finally achieved equilibration with an average of 28.425 ± 0.110 nm during the last 25 ns of the trajectory signifying structural compactness acquired at the conclusion of the simulation.

Next, intramolecular hydrogen bonds were calculated for both wt-hEPO-EPORs and Analog-71.1-EPORs complexes which is another parameter of structural durability. The average H-bonds for wild-type complex and the analog-receptor complex were found to be 153 and 151, respectively. As shown in Figure 9C, there was no significant difference between the trajectory of both complexes, representing conformational steadiness of both complexes.

Subsequent analysis was done on RMSF profile to examine the degree of fluctuation in residue level of the analog after binding to the receptors. Figure 10 demonstrates RMSF of both analog-receptors and hEPO-receptors complexes plotted against residue number. As displayed in the RMSF plot, the highest peak of the analog was for D^165^ (8.179 nm) which is located in the C-terminal region of the molecule. Other peaks were found to be for A^30^, S^84^, and A^125^ which are located in the loop regions of the molecule. Apart from these residues, the pattern of residual oscillation of Analog-71.1 was very similar to that of the wild-type molecule. The in-depth analysis revealed that the residues of the analog interacting with the receptors, which were identified from docking studies (Table 4), did not show any signification variation from their native counterparts during MD simulation, indicating stable interaction of the analog with the receptors. Moreover, the modified region of the analog, marked by black dotted box in the plot, also showed less flexibility. Consequently, the flexibility profile confirmed the stable conformation of Analog-71.1 while interacting with the receptors.

**Figure 10:**
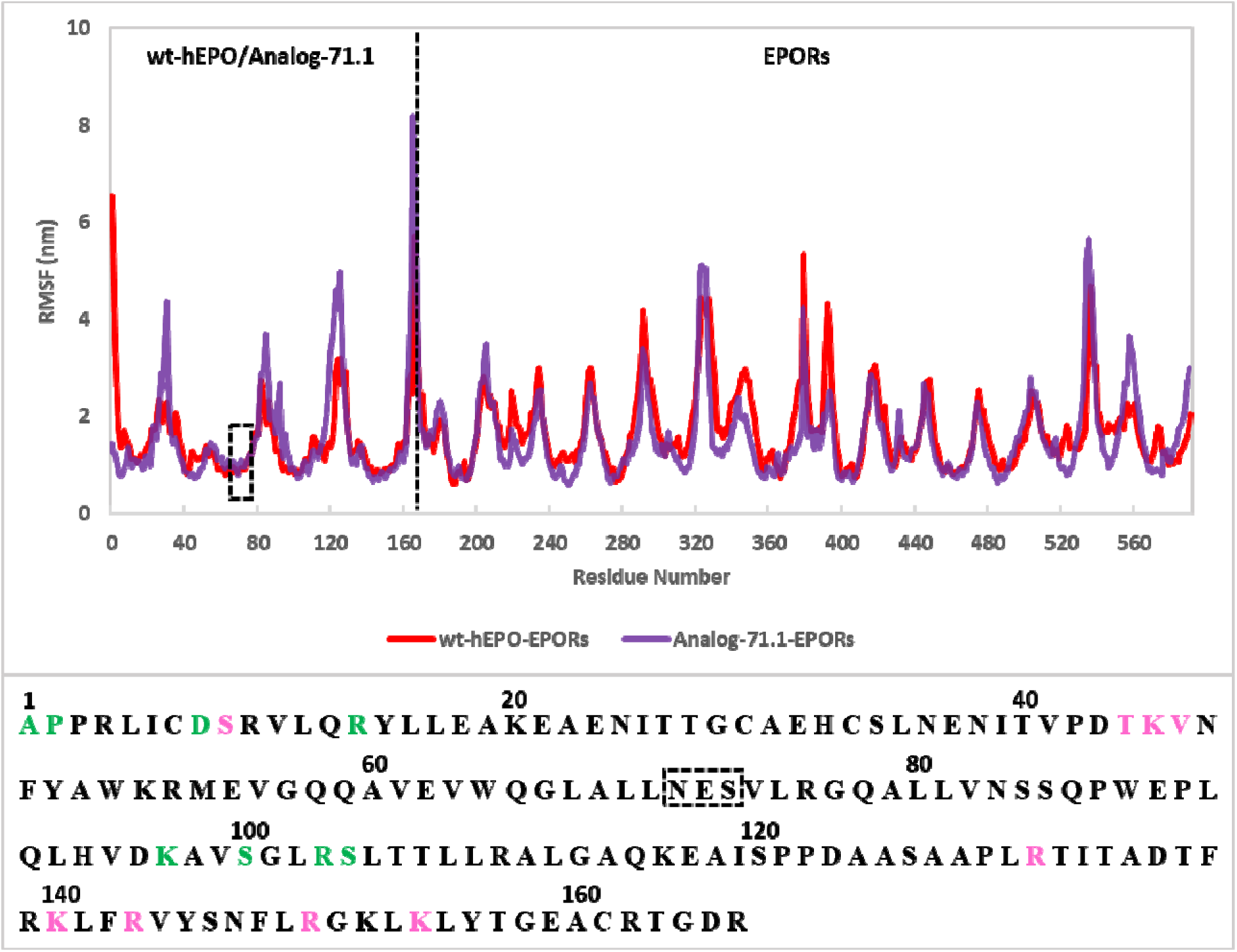
RMSF profile plotted against residue number for Analog-71.1-EPORs complex (purple) in comparison with wt-hEPO-EPORs complex (red) obtained from 100 ns MD simulation. The amino acid sequence of Analog-71.1 is displayed below the RMSF plot where the newly introduced N-glycosylation site is marked by black dotted box and the residues present at the high affinity and low affinity interaction interface are indicated by pink and green colors respectively. No significant fluctuation was observed in the RMSF plot for any of these critically important residues.

## Discussion

Recombinant human erythropoietin (rhEPO) is a crucial glycoprotein drug which is used in the treatment of anemia associated with various clinical conditions (Elliott et al., 2003). To increase the efficiency of any glycoprotein drug, glycoengineering i.e., increasing carbohydrate content of the protein is a common and effective strategy (Shafaghi et al., 2019). Addition of N-glycosylation at appropriate position can improve biological function of glycoprotein therapeutics by increasing molecular stability and serum half-life (Ceaglio et al., 2008; Costa et al., 2014; Sinclair & Elliott, 2005). However, this strategy can also reduce bioactivity by decreasing receptor binding affinity or causing protein misfolding, if carbohydrate moiety is incorporated at wrong position (Samoudi et al., 2015). Therefore, it is important to find out appropriate position for glycosylation. But modifying multiple positions of a glycoprotein in laboratory to find out suitable one(s) for glycosylation is time consuming and expensive (Samoudi et al., 2015). So, a rational *in silico* approach can be highly valuable to shortlist potential positions for glycosylation before going through tedious laboratory work. For example, hyperglycosylated analogs of human interferon beta, human luteinizing hormone and human follicle-stimulation hormone were designed, modeled and analyzed through computational method in various research works conducted recently (Bahadori et al., 2022; Samoudi et al., 2015; Shafaghi et al., 2019). On the basis of this, the aim of our current study was to build computational model of prospective hyperglycosylated analogs of human erythropoietin and evaluate them through molecular docking and molecular dynamics simulation. In a previous study, a hyperglycosylated hEPO analog was designed and modelled, but the interaction of the analog with the receptors and structural stability of the analog in dynamic environment were not explored (Kianmehr et al., 2015).

In our present study, forty mutant analogs were designed by inserting N-glycosylation consensus sequence (N-X-S/T) at different positions in αB helix (55-83 residues) of hEPO through amino acid substitution, since no receptor binding residues are located in this region (Syed et al., 1998). It was expected that carbohydrate addition should not hamper the interaction of the molecule with the receptors. Three dimensional models of the analogs were generated by homology modeling and the probability of their glycosylation was checked according to their sequence context as well as spatial arrangement in 3D conformation. Twenty of forty analogs, which had either glycosylation probability score <50% or spatial inaccessibility for glycosylation or distorted docking structure, were excluded from subsequent analysis (Table 1). The remaining twenty analogs, while evaluating their computational models, showed that majority of their residues belonged to the most favored regions and additional allowed regions in the Ramachandran plot, ensuring good quality of their structures (Supplementary Table 1). Quality score of ERRAT and other evaluating parameters used in this study further validated the acceptability of the mutants’ 3D structures. For *in silico* glycosylation of the hEPO analogs, the structure of carbohydrate chain was determined by studying previous literatures. According to Sasaki and his co-workers (1987), the quantity of tetraantennary saccharides attached to the native hEPO hormone is greater than the other biantennary or triantennary saccharides (Sasaki et al., 1987). They also observed these oligosaccharides to be present mostly in trisialosyl forms (Sasaki et al., 1987). Therefore, a tetraantennary carbohydrate chain containing three terminal sialic acids was selected from the database of GlyProt which was combined at all the four N-glycosylation sites of the analogs to build the hyperglycosylated form of the analogs.

The hyperglycosylated analogs were then subjected to molecular docking in order to examine their interaction with the erythropoietin receptors. The docking scores of the docked complexes are recorded in Table 1. The greater docking scores referred to better pose of the analogs with the receptors. Moreover, to determine which analog(s) would bind to the receptors more strongly and more appropriately, further in-depth analysis was conducted by assessing the number of hydrogen bonds formed within the docked complexes and identifying the residues involved in those bonds. Then MD simulation was performed in two stages. Firstly, the homology models of the selected analogs were simulated for 50 ns to investigate whether the sequence modification for N-glycosylation would cause any distortion in analogs’ structural stability, compactness, flexibility and other important characteristics. Secondly, 100 ns MD simulation was conducted for the best qualified analog-receptor docked complex, identified from molecular docking and 50 ns simulation study, in order to scrutinize the interaction of the analog with the receptors in dynamic environment.

Through docking study, we discovered that all high scoring docked complexes didn’t have higher number of hydrogen bonds. For example, Analog-72.2 obtained the highest docking score (2338.113), but formed moderate number of hydrogen bonds (15) with the receptors. Again, Analog-58.2, one of the top scoring analogs (2111.764), had lowest number of hydrogen bonds (6). So, these analogs were considered as less potential candidates. Subsequently, five analogs were selected for MD simulation which had both high docking scores and higher number of hydrogen bonds.

Among the qualified analogs, Analog-64.2 attained good docking score (2050.646) and formed 20 hydrogen bonds with the receptors. However, the MD simulation study showed large deviation in the RMSD trajectory of the analog, indicating lack of stability of the analog’s structure (Figure 6A). The analog also had highly fluctuating residues in its C-terminal region as shown by its RMSF profile (Figure 7A). The docking score and the number of hydrogen bonds of Analog-72.1 (2051.840, 19) were close to Analog-79.1 (2068.152, 20). Both of these analogs attained equilibration in their RMSD during simulation, but that did not continue till the end of the trajectory (Figure 6C and 6E). At the later phase of the simulation, the RMSD plot of Analog-72.1 inclined upwardly, while Analog-79.1 showed fluctuations. The RMSF plots also displayed higher peaks of residual oscillation for both of these analogs in comparison to wild-type hEPO (Figure 7C and 7E). Analog-76.1-EPORs complex exhibited favorable docking score (2025.016) and very prominent number of hydrogen bonds (24). Out of 24 bonds, the analog formed 9 bonds with B chain and 15 bonds with C chain of the receptors. The pattern of bond formation was unlike the natural hEPO which forms more hydrogen bonds (16) with B chain and less (13) with C chain of the receptors creating higher affinity and lower affinity interaction interface (Table 4). The RMSD plot obtained from 50 ns simulation of Analog-76.1 was comparatively stable than Analog-64.2, Analog-72.1 and Analog-79.1 (Figure 6D). However, highly fluctuating residues were observed in the C-terminal region of Analog-76.1 similar to Analog-64.2 and Analog-79.1 as revealed by RMSF analysis (Figure 7D). The analog also showed less stability in the trajectory of R_gyr_ and SASA (Figure 8B and 8D).

The analysis conducted by molecular docking and MD simulation demonstrated the most promising results for Analog-71.1. According to the docking study, Analog-71.1-EPORs complex had the highest number of hydrogen bonds (25) which also achieved second highest docking score (2313.711), anticipating strong binding affinity of the analog with the receptors. Amid 25 bonds, 13 were formed with EPOR B-chain and 12 with EPOR C-chain, resembling to the balanced interaction of wild-type hEPO-EPORs complex. Besides, the residues of Analog-71.1 involved in receptor binding are mostly similar to the wild-type molecule (Table 4)

The RMSD profile of the analog showed that the equilibrium state was achieved pretty soon during simulation which was continued with a steady trajectory till the end, indicating stable conformation of the analog (Figure 6B). Particularly, after 35 ns the analog maintained constantly lower RMSD value than wt-hEPO. This suggested improved stability of the analog than the wild-type. The R_gyr_ profile supported the result obtained from RMSD analysis by showcasing steady plateau with less fluctuation throughout the simulation ensuring structural compactness. Additional study was conducted through RMSF analysis, the higher value of which represents more flexibility and less stability in protein structure (Borjian Boroujeni et al., 2021). The RMSF plot displayed that the amino acid replacement in the modified region of the analog didn’t cause any notable variation in the residual fluctuation level than wt-hEPO (Figure 7B). However, elevated fluctuation was seen for the residues of position 83-86 in the analog, but these residues fall into loop region, so little fluctuation in this region is acceptable. Solvent accessibility of the analog was also computed through MD simulation and compared with that of wild type molecule. Higher SASA values of a protein usually denotes structural relaxation and thereby decreased stability in protein structure and vice versa (Borjian Boroujeni et al., 2021). In our study, the average SASA value of Analog-71.1 was lower than the wild type which represents another confirmation for analog’s stability (Table 5). Again, the MD simulation of the analog produced a steady trajectory for intermolecular hydrogen bond formation ensuring the stabilized conformation of the analog (Figure 8E).

After checking the stability of Analog-71.1 through 50 ns MD simulation, 100 ns simulation study was employed with the aim to examine the stability of analog-receptors complex structure. Initially the RMSD plot of Analog-71.1-EPORs showed more deviation than the wild-type complex, but eventually equilibration was achieved which was continued till the end of the simulation (Figure 9A). Relatively more fluctuation was seen in the R_gyr_ plot of Analog-71.1-EPORs complex from 22 ns to 76 ns in comparison to wt-hEPO-EPORs (Figure 9B). After that the analog reached the plateau expressing structural compactness earned at the end of the simulation. Furthermore, the stable interaction of Analog-71.1 with the receptors was ensured by studying their RMSF profile which showcased that the level fluctuation of the interacting residues of Analog-71.1 was almost indistinguishable from their native counterparts (Figure 10).

Taking everything into account, Analog-71.1 represents the most promising candidate in our analysis to develop analogous drug of human erythropoietin. Further experiments in the laboratory needs to be conveyed to confirm the efficiency of the analog.

## Conclusion

In today’s date, computational approach in drug discovery is on high demand, since it can shortlist potential drug candidates before going through intensive laboratory work. Since recombinant hEPO is a very valuable drug used in lots of health conditions causing anemia, discovery of its more efficient analogous drug would be of great use. In our study, we attempted a rational *in silico* approach to design and model multiple hyperglycosylated hEPO analogs and identify the best candidate through proper analysis. From our study, it is expected that Analog-71.1 would maintain a stable conformation after sequence modification and it would have strong and stable interaction with the receptor dimer, which made the analog a potential drug candidate for future *in vitro* study.

## Data Availability

All data generated or analyzed during this study are included in this published article.

## Authorship contribution statement

**NT:** Conceptualization, Methodology, Investigation, Data Curation, Formal analysis, Visualization, Writing - Original Draft Preparation, Review & Editing. **KIK:** Investigation. **MNS:** Writing – review & editing. **MSR:** Conceptualization, Supervision, Writing - Review & Editing.

## Acknowledgements

The authors would like to thank Abu Nasor Md. Rakib Sarwar for his valuable technical support in the investigation.

