## Supplementary File for "Unveiling Novel Hyperglycosylated Analog of Human Erythropoietin: A Comprehensive Computational Exploration"

**Supplementary Material**

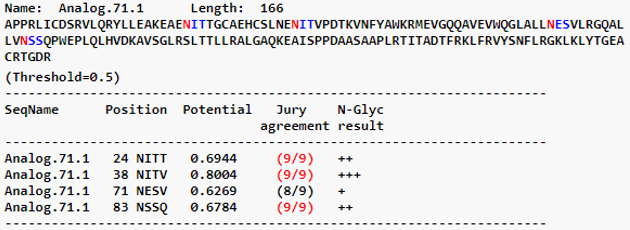

**(A)**

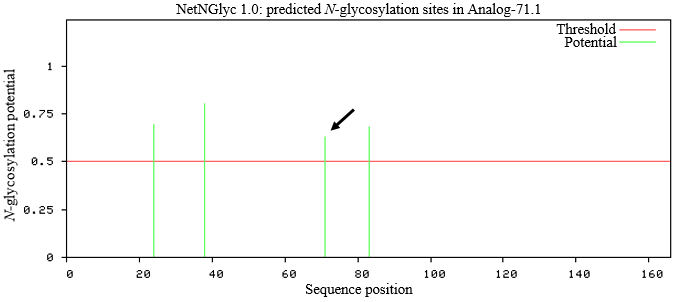

**(B)**

**Supplementary Figure 1:** Predicted N-glycosylation sites in Analog-71.1. (A) The table displays potential scores of four probable N-glycosylation sites of Analog-71.1, all of which obtained positive scores. (B) In the graph, the potential scores for the probable N-glycosylated sites were plotted against sequence position. The vertical green lines represent the asparagines (N^24^, N^38^, N^71^ and N^83^) potential for N-glycosylation crossing the threshold level (0.5), the horizontal red line represents the threshold level and the black arrow marks the newly inserted asparagine (N^71^) residue.

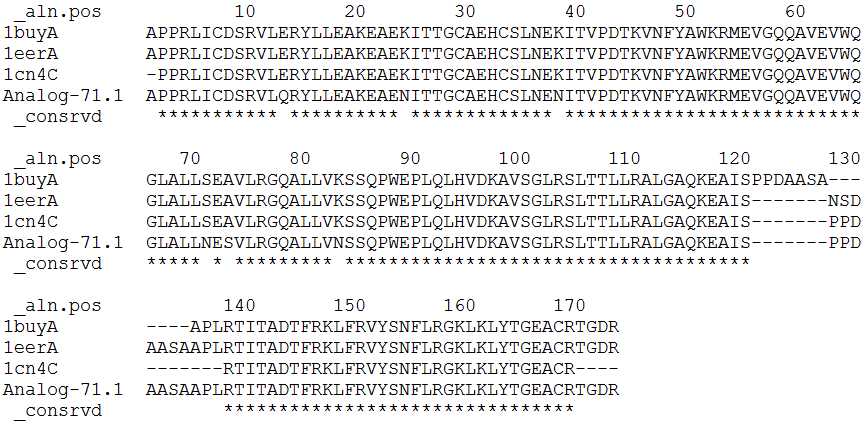

**Supplementary Figure 2:** Multiple sequence alignment of Analog-71.1 with the templates showed that most of the regions of the molecules were identical.

**Supplementary Table 1**: Evaluation parameters of homology models of hEPO analogs.

| Analogs | Ramachandran plot | | | | Quality factor | Whole RMSD (nm) | Z-score |
| --- | --- | --- | --- | --- | --- | --- | --- |
|  | Most favored regions | Additional allowed regions | Generously allowed regions | Disallowed regions |  |  |  |
| Analog-55.1 | 89.1% | 9.5% | 0.7% | 0.7% | 85.4430 | 1.32 | -6.62 |
| Analog-55.2 | 91.2% | 7.4% | 1.4% | 0.0% | 86.6242 | 1.41 | -6.74 |
| Analog-56.1 | 89.1% | 9.5% | 0.7% | 0.7% | 85.4430 | 1.38 | -6.69 |
| Analog-56.2 | 92.5% | 6.1% | 1.4% | 0.0% | 90.3226 | 1.35 | -6.72 |
| Analog-57.1 | 91.2% | 6.8% | 2.0% | 0.0% | 87.3418 | 1.02 | -6.05 |
| Analog-57.2 | 91.2% | 7.4% | 1.4% | 0.0% | 87.3418 | 1.46 | -6.75 |
| Analog-58.1 | 95.2% | 4.1% | 0.7% | 0.0% | 94.9367 | 1.40 | -6.02 |
| Analog-58.2 | 93.9% | 6.1% | 0.0% | 0.0% | 82.2785 | 1.28 | -6.13 |
| Analog-64.2 | 94.6% | 5.4% | 0.0% | 0.0% | 84.0764 | 1.26 | -5.93 |
| Analog-69.1 | 93.9% | 3.4% | 2.0% | 0.7% | 98.1013 | 1.28 | -6.24 |
| Analog-69.2 | 93.9% | 5.4% | 0.7% | 0.0% | 89.7436 | 1.06 | -6.20 |
| Analog-71.1 | 92.5% | 6.8% | 0.7% | 0.0% | 96.2025 | 1.27 | -6.48 |
| Analog-71.2 | 94.6% | 4.8% | 0.7% | 0.0% | 91.1392 | 1.60 | -5.99 |
| Analog-72.1 | 93.2% | 6.8% | 0.0% | 0.0% | 89.8734 | 1.19 | -5.96 |
| Analog-72.2 | 91.2% | 7.5% | 0.7% | 0.7% | 85.4430 | 0.84 | -5.92 |
| Analog-75.1 | 93.2% | 6.8% | 0.0% | 0.0% | 80.3797 | 1.18 | -6.06 |
| Analog-75.2 | 94.6% | 5.4% | 0.0% | 0.0% | 94.9045 | 1.31 | -6.55 |
| Analog-76.1 | 92.5% | 6.8% | 0.0% | 0.7% | 91.7722 | 1.63 | -6.13 |
| Analog-76.2 | 90.5% | 9.5% | 0.0% | 0.0% | 79.1139 | 1.37 | -6.01 |
| Analog-79.1 | 93.2% | 5.4% | 0.7% | 0.7% | 83.5443 | 1.59 | -6.30 |

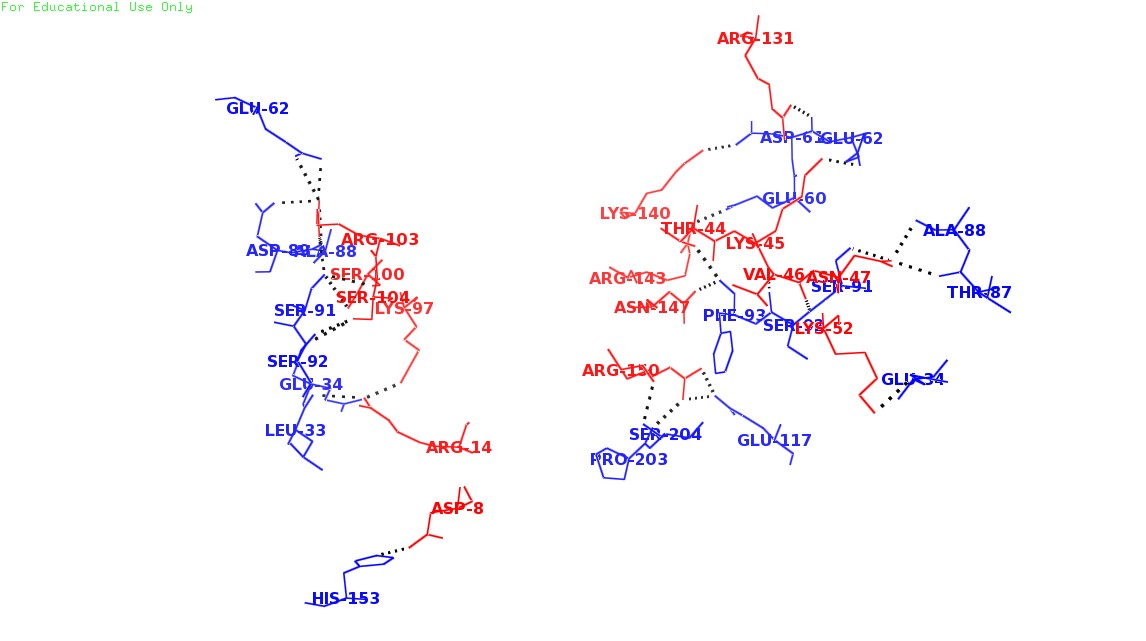

**Supplementary Figure 3:** Interaction interface between wild-type hEPO and EPORs; red colored lines representing amino acids of wt-hEPO and blue colored lines representing amino acids of the receptors**,** black dots representing the hydrogen bonds formed among the amino acids.
